# Auditory features modelling reveals sound envelope representation in striate cortex

**DOI:** 10.1101/2020.04.15.043174

**Authors:** Alice Martinelli, Giacomo Handjaras, Monica Betta, Andrea Leo, Luca Cecchetti, Pietro Pietrini, Emiliano Ricciardi, Davide Bottari

## Abstract

The striate cortex is no longer considered exclusively visual in its function. Proofs that its activity is modulated by acoustic inputs have accrued. By employing category-based and feature modeling approaches, here we characterized V1 activity (in absence of retinal input) during the processing of natural and synthetically derived sounds. First, we showed that distinct sound categories could be dissociated by the analysis of V1 multivoxel response patterns. Hence, we assessed whether a hallmark of sound neural representations is mapped in V1. In each sound category, we modeled sound envelopes and assessed whether these were represented at the single-voxel level in the striate cortex and, as a control, in the temporal cortex. The hierarchical organization of sound categories allowed to exert control over dimensions that could spuriously lead to sound envelope V1 mapping. Variations of sound amplitude over time were successfully decoded in V1 regardless of the category class. Results confirm that the human striate cortex receives acoustic category-based input and demonstrate that V1 is a genuine locus of sound envelope representation.

## Introduction

Perception has been considered as highly segregated for more than a century. The existence of specialized detectors for different forms of environmental energies has prompted the enduring dominant paradigm, which postulates that information from different sensory modalities is anatomo-functionally segregated in the human brain (Pascual-Leone and Hamilton, 2001; Scholvinck et al., 2010). This assumption received strong supporting evidence from seminal lesion studies demonstrating that unimodal deficits were associated with lesions in primary sensory cortices (Massopust et al., 1965; Winans, 1967). In particular, the visual system has been thought to be functionally independent of cross-modal influences and, if anything, to dominate the other senses (McGurk, 1976; Pick et al., 1969).

With the advent of multisensory research (Stein and Meredith, 1993), the last decades have witnessed the challenging of such traditional view. A burgeoning of animal and human studies proposed that cortical areas, previously believed to only process sensory-specific information, would be multisensory instead (Ghazanfar and Schroeder, 2006; Schroeder and Foxe, 2005).

In humans, the corroboration that, at least to some degree, multisensory interplay occurs already at the level of V1 has been reported using hemodynamic imaging (Eckert et al., 2008; Martuzzi et al., 2007; Rohe and Noppeney, 2016; Zangenehpour and Zatorre, 2010) and electrophysiological measures (Giard and Peronnet, 1999; Mercier et al., 2013; for a review see Murray et al., 2016). To challenge even more the modality-segregated view, evidence of pure cross-modal responses in V1, evoked by unimodal auditory inputs, exists. The modulation of the oscillatory activity in the striate cortex has been observed in response to pure tones (Mercier et al., 2013; Romei et al., 2012). Sound-driven responses in V1 have also been measured with intracranial recordings: high gamma neural oscillations after white-noise bursts were measured in the striate cortex (Ferraro et al., 2020).

Not all sounds seem to exert the same effect on V1. High pitch and narrowband sounds elicit a more significant increase of visual cortex excitability compared to lower pitch and broadband sounds, respectively (Spierer et al., 2013). Moreover, looming sounds have been found more effective than static or receding sounds in enhancing visual cortex excitability (Romei et al., 2009). These results revealed stimulus-selective cross-modal interactions in low-level visual cortex. However, which sound features are mapped in human striate cortex is still unknown. All previous works on this issue assessed stimulus-evoked responses by comparing different natural or artificial sound categories (Vetter et al., 2016, 2020; e.g., Spierer et al., 2013; Romei et al., 2009), artificial sounds and silence (Ferraro et al., 2020; Mercier et al., 2013), or by measuring the hemodynamic activity associated to artificial sounds (e.g., Martuzzi et al., 2007). These approaches limit the current understanding of auditory activity in V1 in three aspects: (i) contrasts between sound categories are over-simplifications as they depend on broad stimulus descriptions, varying across multiple experimental dimensions (e.g., visual, acoustic, semantic) and do not allow extracting the fine-grained neural representation of acoustic features. (ii) Responses to simple artificial sounds, such as pure tones, do not allow to exploit the richness of the population encoding properties. Finally, (iii), these approaches do not clarify whether sounds could be encoded in V1 with shared responses across large patches of cortex, similarly to category-based activity patterns (Haxby et al., 2011) or with specific functional units showing a lower-level feature-based tuning (Carandini et al., 2005).

To fill these gaps, following a category-based analysis, we combined auditory modeling of different natural (or synthetically derived) sound categories (i.e., speech, speech-related and non-speech naturalistic sounds) with the measure of brain activity in the absence of retinal input. Natural sounds and vocalizations are characterized by profiles of high power at slow temporal amplitude modulations (AM). Studies in animal models revealed that the statistical structure of natural sounds, such as their characteristic intensity fluctuations, matches the auditory system’s neural coding selectivity (Hsu et al., 2004; Riecke, 1995). Thus, we modelled the envelope power of sounds and specifically assessed whether this hallmark of natural sounds processing is mapped in V1.

Sound amplitude changes occurring at typical speech envelope time-scales were exploited (Di Liberto et al., 2015; Giraud and Poeppel, 2012; Luo and Poeppel, 2007). Four categories of hierarchically organized acoustic stimuli were tested in isolation to maximally characterize population encoding properties (Figure 1B). All of them comprised stimuli with natural amplitude modulations. First, we measured the brain response to selected single words pertaining to the same semantic class. As control stimuli, from each word an associated pseudoword was generated. Pseudowords retained similar articulatory patterns and envelopes of each word from which they were derived, but did not convey any semantic content; these stimuli allowed to control for an association between V1 responses and semantic information (Figure 1A). As a further control condition, for each word stimulus, an artificial sound was also created by preserving the word sound envelope but flattening the original spectral structure (Figure 1A, B). Thus, each artificial sound shared with the originating word only the information provided by its amplitude modulation. This control condition was conceived to isolate the role of sound amplitude modulations and to rule out the contribution of spectral properties occurring in speech, such as the fine harmonic structure. Therefore, we had an associated pseudoword and an artificial sound, with comparable syllabic and phonemic timescales of signal amplitude changes starting from a specific word. Finally, we introduced bird chirps as an additional control condition (Figure 1B). This category allowed to control for speech specificity of the neural responses.

Evoked hemodynamic activity was acquired during a listening task. By employing multivoxel pattern analysis (MVPA), we provided a description of the representational spaces obtained from overall activity of the visual cortex across the four sound categories. Voxel-wise decoding, based on a parsimonious auditory model, evaluated within each sound category whether envelope variations in the frequency ranges of interest were associated to activity of the striate cortex, thus ascribing its response to a fundamental acoustic feature.

**Figure 1.**
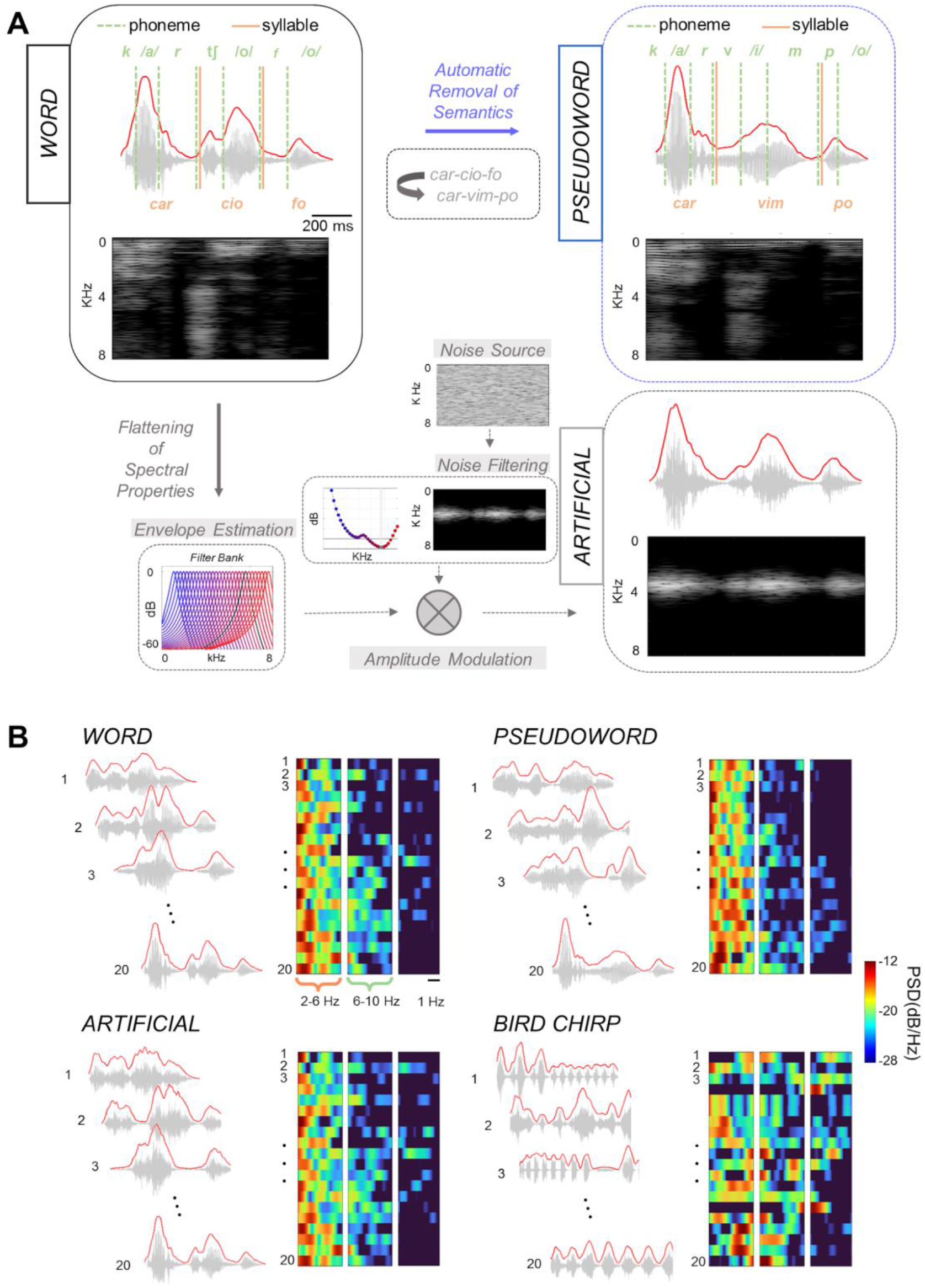
Stimuli. (A) The word sound category comprised 20 tri-/four-syllabic words (mean duration 1.00 ± 0.06 sec) pertaining to the same semantic category (i.e., vegetables). From each word two control stimuli were created: (i) a pseudoword was obtained by using an automatic algorithm. A multilingual pseudoword generator able to produce polysyllabic stimuli that respect the phonotactic constraints of the Italian language was adopted. (ii) An artificial sound was created by preserving the sound envelope of the word of origin but flattening its spectral structure. The original audio trace of each word was decomposed into 30 critical bands, using a gammatone filter-bank. Single sub-band envelopes were then linearly summed across these critical bands and then used to modulate the amplitude of a white band-passed Gaussian noise source whose central frequency was characterized by the lowest absolute threshold of hearing (f = 3.4 kHz). (B) For each stimulus of the four sound categories (word, pseudoword, artificial sounds, and bird chirps) the envelope Power Spectral Density (PSD) was extracted. We defined two bins of interest between 2-6 and 6-10 Hz, representing the modulation power in a Low and High-frequency ranges which were associated with phonemic and syllabic frequencies rates respectively, in speech-related categories. Note that speech-related categories retained only minimal energy above 10 Hz.

## Results

### Activity of the Calcarine cortex across sound categories

Provided the aim of the present study, all the analyses were conducted selectively within two regions of interest (ROIs): (i) the *Occipital ROI* selectively included the Calcarine cortex (i.e., V1) using the probabilistic map by Wang and colleagues (Wang et al., 2015); (ii) the *Temporal ROI* comprised the Superior Temporal Gyrus and Sulcus as well as the Middle Temporal Gyrus (AICHA atlas, (Joliot et al., 2015).

Firstly, we measured Calcarine hemodynamic activity elicited by words, pseudowords, artificial sounds and bird chirps in each participant. Results showed an overall positive response across participants for each sound category (~0.4%, all conditions with p < 0.001, two-tailed, Student’s t-test, Figure 2A), indicating that the early visual cortex was not globally suppressed, but was instead significantly engaged during auditory processing with eyes closed.

Given this evidence, we analyzed the response patterns of the bilateral Occipital ROI by means of a group-level Principal Component Analysis (PCA). The technique decomposed the overall activity elicited by thousands of voxels in 63 PCs explaining 95% of the variance, which were hyper-aligned across individuals (Haxby et al., 2011). From each PC, we obtained the associated Representational Space (RS) and measured the distances between stimuli pertaining to each class compared to distances of stimuli between classes. Results indicated that several PCs retained a category-based representation (i.e., within-category distances were significantly lower than between-category distances, q < 0.01, permutation test, FDR corrected; Figure 2B). We also averaged the RSs generated from each significant PC (Figure 2C) to show that category-based representations are retrieved from activity of the Calcarine cortex. Results confirmed previous studies employing fMRI and MVPA (Vetter et al., 2016, 2020). Nonetheless, category-based representations could be driven by factors, such as imagery or semantics, rather than by acoustic features. Thus, we adopted for subsequent analyses a within-category approach, aiming at reconstructing envelope variations in the frequency ranges of interest for each stimulus, using activity of the Calcarine cortex (and Temporal ROI) as predictor.

**Figure 2.**
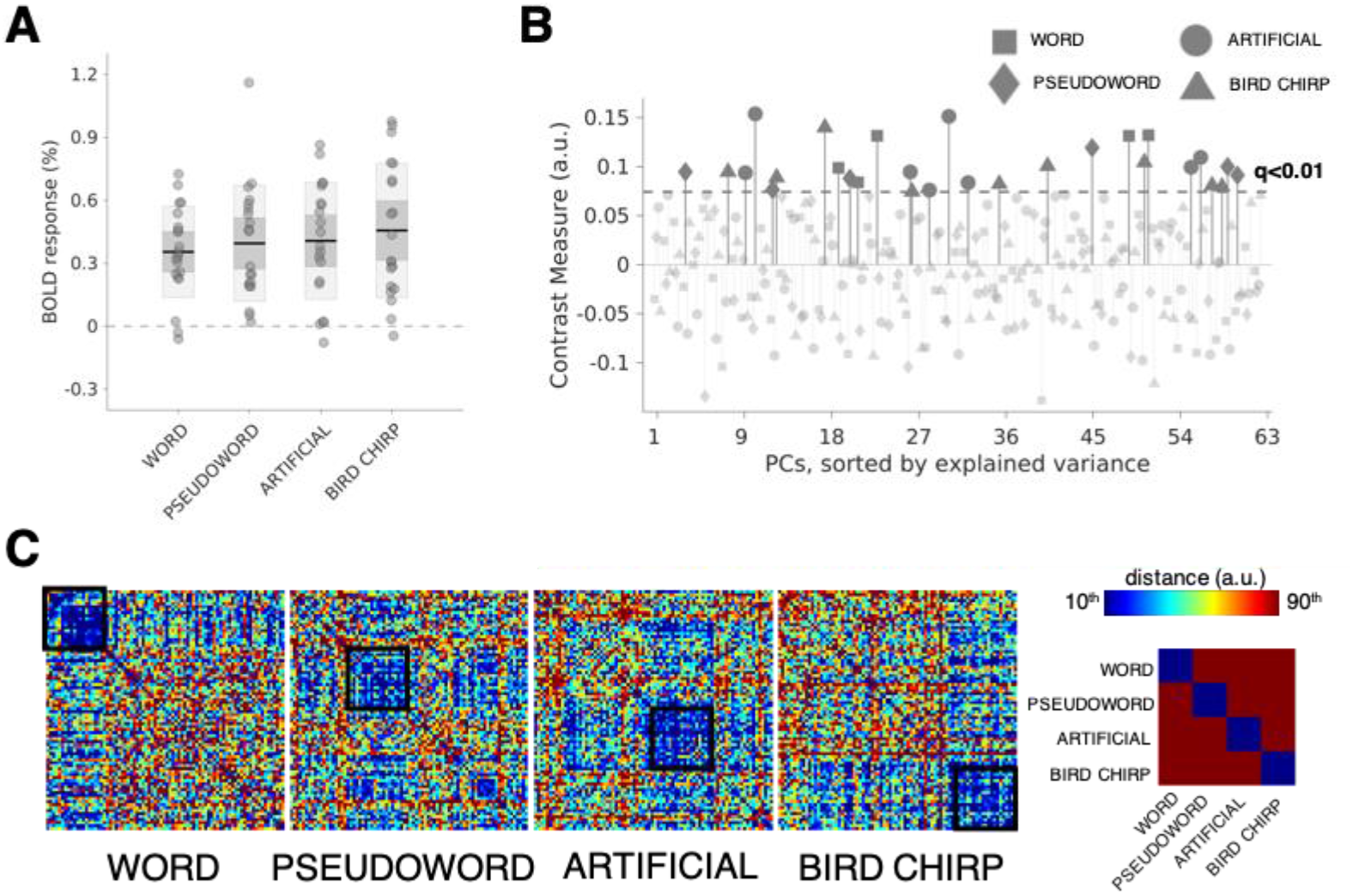
Striate cortex activity across sound categories. (A) fMRI average activity was extracted from striate cortex defined by the probabilistic atlas of Wang and colleagues. All the four sound categories retained a significant positive response (p < 0.001, two-tailed, Student’s t-test). Each dot indicates a participant, the black line within each boxplot represents the mean hemodynamic activity, whereas the dark-gray box indicates its standard error, and the light-gray box represents the standard deviation. (B) We explored activity patterns of the striate cortex by extracting group-level Principal Components (PCs), up to 95% of total explained variance. From each PC and each experimental category, we obtained a measure of contrast, by comparing the Euclidean distances between and within sound categories, where positive values indicated lower within-category distance (i.e., high contrast). The Manhattan plot reports the contrast measures across the four different categories and PCs, sorted in descending order of explained variance. The dotted line indicates the significance level (permutation test; q < 0.01, FDR corrected). Several PCs retained a category-based representation, and their averaged representational spaces were depicted separately for each class in panel (C), using the Euclidean distance and a color scale ranging from 10^th^ to 90^th^ percentile.

### Reconstructing envelope variations from activity of Calcarine and Temporal ROIs

First, we manually measured the syllabic and phonemic frequencies of words and pseudowords (median word syllabic 3.80 Hz, CI-95^th^: 3.62-4.10; word phonemic 8.38 Hz, CI-95^th^: 7.48-9.12; pseudoword syllabic 3.80 Hz, CI-95^th^: 3.54-4.05; pseudoword phonemic 8.37 Hz, CI-95^th^: 7.47-9.67). As previously showed (Keitel et al., 2018), these results were congruent with the envelope power spectral density (PSD) calculated from sound waves (Figure 1B). Considering the dynamic frequency ranges of our linguistic stimuli, in the envelope PSD we defined two non-overlapping 4Hz-wide frequency ranges which were centered, on median values, at 4 and 8 Hz for syllabic and phonemic rates, respectively. From now on, these intervals are identified as Low and High envelope frequency ranges. Note that, as shown in Figure 1B, signals did not comprise substantial energy above 10 Hz. Each artificial sound maintained the sound envelope of the word from which it was generated, but its spectral structure was flattened (Figure 1A). Although bird chirps comprised higher amplitude modulated frequencies as compared to the human speech envelope, for consistency of analyses, their envelope PSD was extracted in the same frequency ranges of words and pseudowords.

**Figure 3.**
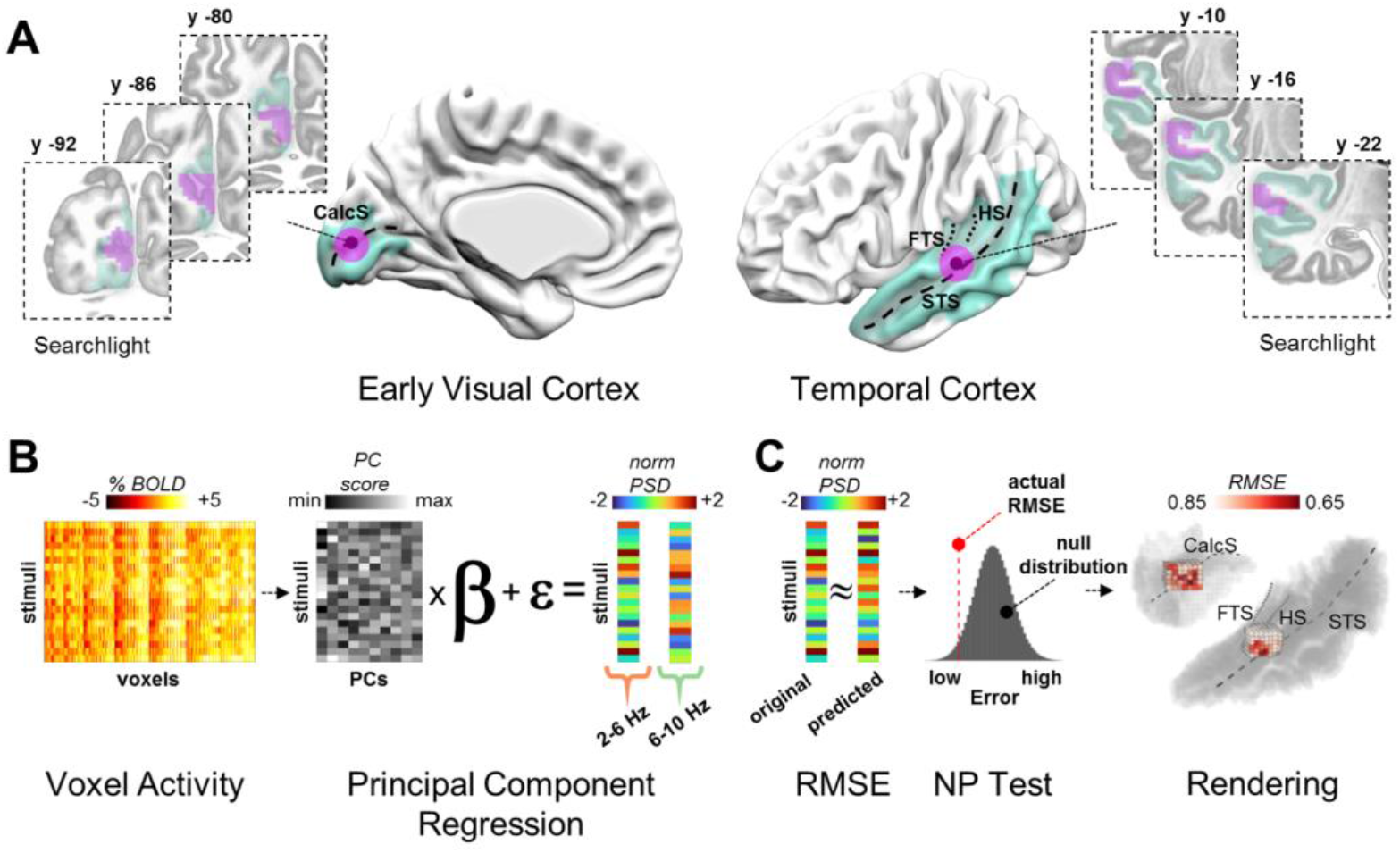
Analysis strategy. (A) fMRI analyses were carried out in temporal and occipital cortical areas. Temporal regions were defined using the AICHA atlas selecting the bilateral Superior Temporal Gyri and Sulci as well as the Middle Temporal Gyri. For the occipital regions, we isolated the striate Calcarine cortex by means of the probabilistic map by Wang and colleagues. A searchlight analysis was performed in the volumetric space respecting the cortical folding. Voxels were sampled along the cortical ribbon preserving the functional distinctions across adjacent sulcal walls and sulci. Calc S indicates the Calcarine Sulcus; FTS indicates First Transversal Sulcus; HS indicates Heschl’s Sulcus; STS indicates the Superior Temporal Sulcus. FTS and HS define the anterior and posterior bounds of Heschl’s Gyrus. (B) Hemodynamic responses were associated with the envelope modulation power using a machine learning algorithm. In each searchlight, we performed a Principal Components (PC) Analysis to extract orthogonal dimensions from normalized voxel responses within each experimental condition and subject. Afterward, we performed a multiple regression analysis, using the PC scores as the independent variable and the power of Low (e.g., 2-6Hz) and High (e.g., 6-10Hz) modulation frequencies as the dependent one. This procedure led to a reconstructed model of predicted power spectral density values in each experimental condition, subject and searchlight. (C) The reconstructed models were compared to the actual ones by calculating their root mean squared error (RMSE). To measure the statistical significance, we first averaged the reconstructed models across participants in each experimental condition and searchlight obtaining a group-level predicted set of acoustic features. Then, we performed a non-parametric (NP) permutation test, by shuffling the predictor matrix (i.e., PC scores) in each subject and experimental condition. This procedure ultimately provided a null distribution of group-level RMSE coefficients, against which the actual association was tested. Results were corrected for multiple comparisons using False Discovery Rate (FDR) and were mapped onto a 3D render of the temporal and occipital regions of interest.

We then reconstructed acoustic features of our stimuli (i.e., the power of the envelope in the High and Low-frequency ranges) by using brain activity as predictor (Figure 3B; Pasley et al., 2012) and mapped the voxelwise root mean squared prediction error (RMSE; Poldrack et al., 2019), after correcting results for multiple comparisons (False Discovery Rate q < 0.01; additional cluster correction of 20 voxels, nearest neighbor). Importantly, to rule out the possibility that whole-brain hemodynamic fluctuations related to changes in arousal and to physiological noise could explain decoding results, we regressed out the global activity defined as the averaged brain response (Aguirre, 1998; Macey et al., 2004). Also, to detail sub-regions of the Calcarine cortex associated with envelope modulations, we mapped the unthresholded RMSE (i.e. below p < 0.01) onto a three-dimensional representation of the region of interest (Figure 3C).

**Figure 4.**
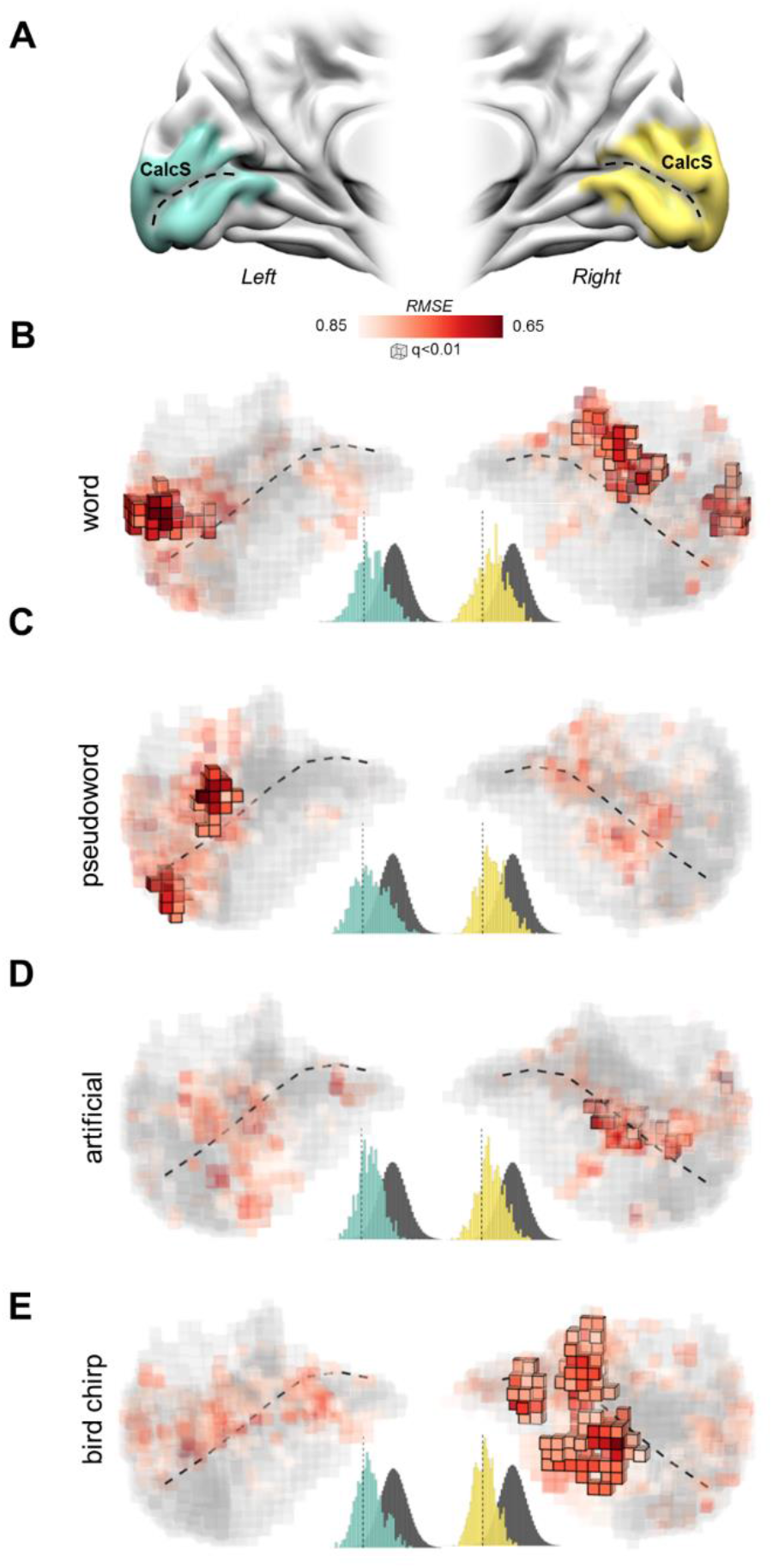
Reconstruction of the sound envelope power in the 6-10 frequency range in V1. (A) V1 ROI in the left (cyan) and right (yellow) hemispheres. (B, C, D, E) Reconstruction in V1 of the sound envelope power in the High (6-10 Hz) frequency range as a function of sound categories (i.e. word, pseudoword, artificial and bird chirps). Significant voxels surviving the multiple comparisons correction are represented by highlighted cubes (q < 0.01; minimum cluster size = 20 voxels), whereas colored cubes displayed uncorrected threshold results at p<0.01. Population histograms represent the overall reconstruction performance in the left (i.e., cyan histogram) and right (i.e., yellow histogram) hemispheres for each ROI and sound category as compared to their null distributions (i.e., dark grey histogram; dashed vertical lines in histograms represents the 1^st^ percentile). Results show that V1 represents sound envelope power for all sound categories with a lower RMSE as compared to the null distribution. The right hemisphere was increasingly engaged from speech stimuli to non-linguistic sounds. For single subject results, see Supplementary Figures 5 and 7. For anatomical landmarks please refer to Figure 3.

The envelope power measured in the High-frequency range, namely the phonemic occurrence in our linguistic stimuli, represented the fastest timescale of the sound amplitude variations we investigated. In a hierarchical feedforward processing scheme the coding of higher frequencies would occur prior to the coding of lower frequencies (DeWitt and Rauschecker, 2012). Such conceptual framework has been suggested for both auditory and visual systems (DeWitt and Rauschecker, 2012; Hubel and Wiesel, 1962). Thus, results that emerged by assessing brain activity associated with the envelope power in the High-frequency range represented the focus of the analysis. Note that comparable results emerged for the two frequency ranges we investigated. The details concerning results associated with the Low-frequency range model (representing the syllabic rate) can be consulted in the Supplementary materials (Supplementary Figure 1, 2, 6 and 8).

### Testing the activity associated with word listening

On these premises, for each word we assessed whether the variation of the envelope power in the phonemic range was associated with the activity of V1 voxels. Results revealed a significant cluster in the posterior part of the left Calcarine sulcus which comprised ~2.8% of the total volume of V1. Moreover, in the right hemisphere, ~4.6% of voxels survived to multiple comparisons correction, distributed across two patches of cortex, one in the middle of the Calcarine sulcus, and an additional smaller cluster in the lateral portion of the Cuneus (Figure 4A). The comparison between the number of significant voxels of V1 in the left and right hemispheres did not differ (χ^2^ _(1, 334)_ = 0.631, p = 0.427; degrees of freedom were estimated by means of Montecarlo procedure, see Materials and Methods). Population histograms further emphasized the overall ability to reconstruct envelope power variations in the High-frequency range, in both left and right V1 as compared to a null distribution.

In line with previous evidence (DeWitt and Rauschecker, 2012) revealing the key role of temporal areas in speech processing, the envelope power in the High-frequency range was successfully reconstructed in these regions. As expected, envelope variations were mainly linked to the brain activity measured in the left hemisphere as compared to the right (~2.7% of the total volume of the left Temporal ROI; ~0.8% of the total volume of the right Temporal ROI; number of voxels in left vs. right hemisphere: χ^2^_(1, 1615)_ = 9.304, p = 0.002). Significant patches of cortex were identified in the left posterior part of the Superior Temporal Sulcus and Gyrus (pSTS/pSTG), which are pivotal regions in phonemic processing (DeWitt and Rauschecker, 2012), as well as in the left mid portions of Middle Temporal Gyrus (mMTG) and in the right central portion of STS (mSTS) (Figure 5B).

The same analysis was performed for the variation of the envelope power in the Low frequency range. The pattern of results was overlapping with previous one (see Supplementary Figure 1; see Supplementary Figure 2 for results in the Temporal ROI). Overall, these results suggest that features extracted from the sound envelope of single words were traceable not only in temporal areas but in the Calcarine cortex as well. However, imagery processes, which are known to activate V1 (Cichy et al., 2012; Naselaris et al., 2015; Vetter et al., 2014) could explain such activations. All word stimuli pertained to the same concrete semantic class favoring coherent imageability across participants.

**Figure 5.**
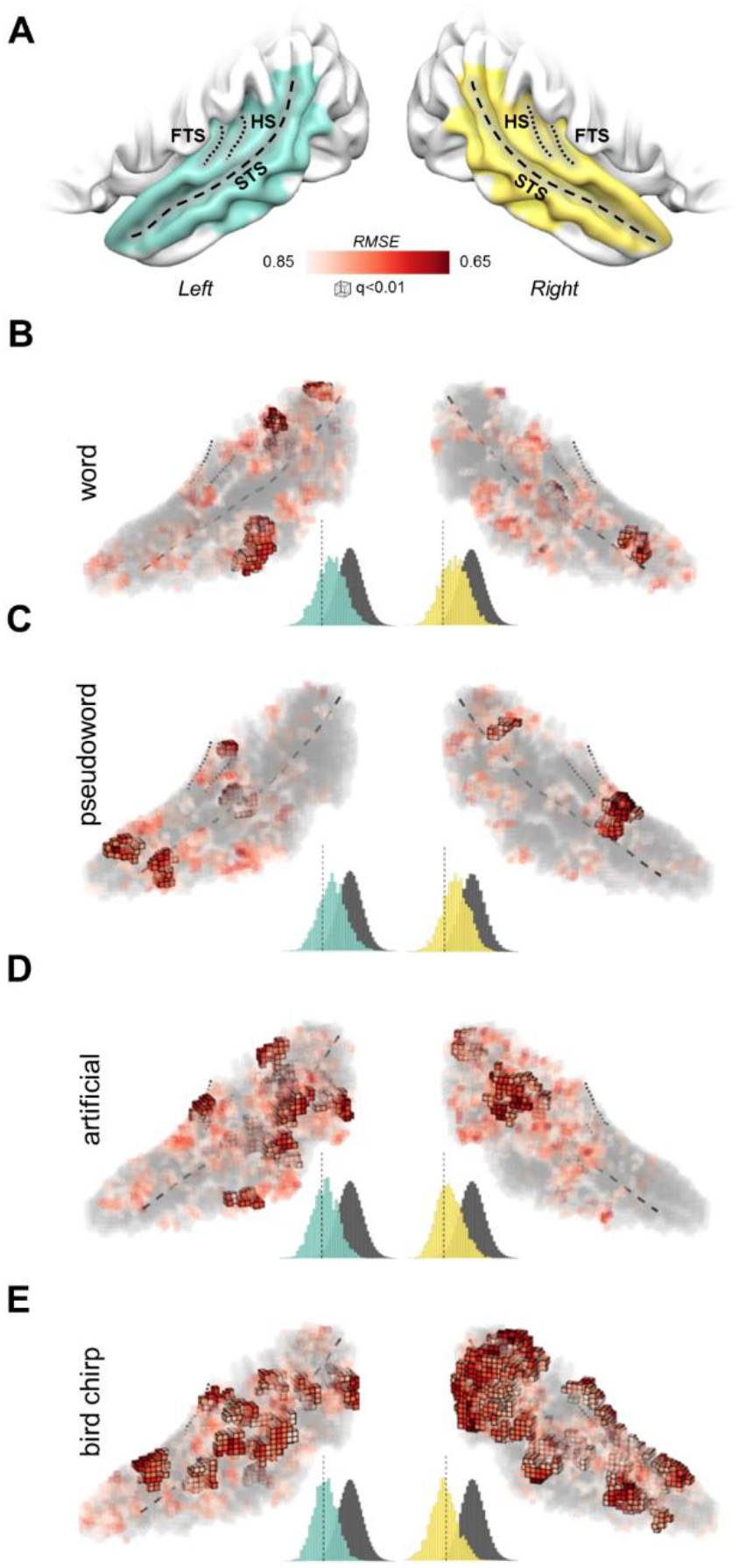
Reconstruction of the sound envelope power in the 6-10 Hz frequency range in the Temporal ROI. (A) Temporal ROI in the left (cyan) and right (yellow) hemispheres. (B, C, D, E) Reconstruction in the temporal ROI of the sound envelope power in the High (6-10 Hz) frequency range as a function of sound categories (i.e. word, pseudoword, artificial and bird chirps). Significant voxels surviving the multiple testing correction (q<0.01; minimum cluster size = 20 voxels) are represented by highlighted cubes, whereas colored cubes displayed uncorrected threshold results at p<0.01. Population histograms represent the overall reconstruction performance for each ROI and sound category as compared to null distributions. Significant clusters emerged in Superior Temporal Sulcus and Gyrus (pSTS/pSTG) in both left and right hemispheres. For anatomical landmarks please refer to Figure 3.

### Testing the role of semantic content

Does the representation of sound envelope in the Calcarine cortex depend on semantic processing and imageability of words? To answer this question, we tested whether the association between brain activity and envelope power variations was still present in the absence of semantic information. Therefore, we measured the brain response to meaningless pseudowords, that retained similar articulatory patterns and phonotactic constraints of the original words from which they were derived (see Figure 1A).

Results in the *Occipital ROI* were consistent between pseudowords and words. Envelope variations in the High-frequency range were associated with two significant clusters in the posterior part of the left Calcarine cortex, extending from the superior bank of the Calcarine sulcus to the Cuneus, and from the inferior bank to the Lingual Gyrus, respectively. Overall ~3.2% of the total volume of left V1 was engaged, whereas no voxels survived to multiple comparisons correction in the right hemisphere (Figure 4C). The number of left V1 voxels associated with the envelope power of pseudowords in the High-frequency range was statistically greater than right V1 (χ^2^_(1, 334)_ = 5.226, p = 0.022; see Figure 4C).

As expected, variations of the envelope power in the High frequency range were also successfully reconstructed in the *Temporal ROI*. Specifically, ~1.9% of the voxels in the left hemisphere and ~1.5% in the right hemisphere were significantly modulated by the envelope in the phonemic range. This revealed the lack of hemispheric dominance for the representation of the envelope power of pseudowords in the High-frequency range (χ^2^_(1, 1615)_ = 0.339, p = 0.560). Significant patches of temporal cortex were identified in the left anterior part of the Superior Temporal Sulcus and Gyrus (aSTS/aSTG), as well as the left mMTG and in the right central portion of STG (mSTG) and pSTS (Figure 5C).

Significant clusters in the Calcarine cortex were found for variations of the envelope power in the Low-frequency range as well (corresponding to the syllabic rate; see Supplementary Figure 1), suggesting that the modulation of V1 activity was not selective for the 6-10 Hz frequency range (see Supplementary Figure 2 for results in the Temporal ROI). Taken together, these findings demonstrate that features of the envelope modulation are represented in the Calcarine cortex even in the absence of semantic content.

### Testing the role of spectral properties

Sound amplitude modulation conveyed by envelope power variations in both Low and High-frequency ranges is known to be associated with changes in sound spectral properties (Di Liberto et al., 2015).

Do the observed results depend specifically on amplitude modulations? Do they rather rely on spectral properties of speech sounds? Or, perhaps, on their combination? Speech comprises envelope variations with specific higher frequency spectral features (e.g., Santoro et al., 2017). Thus, to assess the specificity of the information conveyed by envelope variations, we measured brain responses to additional control stimuli. For each word, we generated an artificial sound of the same duration and estimated envelope, removing the original spectral properties. Specifically, white noise was filtered within the most sensitive hearing frequency band (3.4 kHz) and modulated according to the envelope of the original word (see Figure 1A). We then tested the intelligibility of artificial stimuli by means of an additional control experiment. Participants listened to each artificial sound and were then asked to identify the original sound from which it was generated by means of a two-alternative forced-choice task. For an artificial sound derived from a word, participants had to identify the original sound choosing between the word and its associated pseudoword. Results demonstrated chance level accuracy (mean accuracy ± SE: 51.5% ± 3%, t(9) = 0.52, p = 0.307, one-tailed Student’s t-test, see Supplementary Figure 4B), indicating that intelligibility was not retained despite envelope modulations were maintained. Results of the fMRI experiment using artificial sounds highlighted a cluster in the mid-posterior part of the sulcus in the right Calcarine cortex (~1.9%) and no involvement of its left homologue (left = 0%; Figure 4D). The comparison between the number of voxels of V1 associated with the envelope power of artificial sounds in the High-frequency range did not differ between the left and right hemisphere (χ^2^_(1, 334)_ = 2.938, p = 0.086).

The envelope power in the High-frequency range was also successfully reconstructed in the temporal cortex. A larger percentage of voxels in the left hemisphere were significantly modulated by the envelope power compared to the right hemisphere (left hemisphere ~7.2%, right hemisphere ~2.6%, χ^2^_(1, 1615)_ = 18.22, p < 0.001). Significant patches of cortex were identified in the bilateral pSTS/pSTG, the left mSTS/mSTG as well as the left Heschl Gyrus (HG; Figure 4D).

Comparable results were found in the Low-frequency range: the mid-posterior part of the right Calcarine sulcus was found to significantly encode envelope power variations corresponding to the syllabic rate (please see Supplementary Figure 1 and Supplementary Figure 2 for details).

By analyzing brain activity associated to artificial sounds, we showed that even in the absence of spectral properties, the amplitude modulation of a sound (in both frequency ranges tested here) is represented in the early visual cortex. These results suggested that fine spectral details associated with envelope variations could not account for the specific response in V1. Overall, these findings support the central role of sound envelope in modulating occipital (and temporal) activity. However, they do not rule out the possibility that specific speech-like temporal modulation of the envelope may be responsible for such an effect.

### Testing the speech specificity

Does Calcarine activity encode envelope variations of non-speech natural sounds as well? To determine whether there is a speech specificity for the observed effects, we examined brain responses to natural sounds characterized by strong envelope power modulation other than human utterances: bird chirps.

Significant clusters associated with the High envelope variation frequency range were found in the middle part of the right, but not of the left, Calcarine cortex (Figure 4E). Specifically, significant patches comprised both the superior and inferior bank of the right Calcarine sulcus. Results revealed the engagement of ~9.4% of the total volume of right V1, whereas left V1 was not involved (right > left: χ^2^_(1, 334)_ = 16.304, p < 0.001).

As expected, the envelope power in the High-frequency range was reconstructed in the temporal cortex as well. Specifically, a large extent of cortex successfully retained envelope characteristics: ~8.2% of the voxels in the left hemisphere and ~21.5% in the right hemisphere (χ^2^_(1, 1615)_ = 57.077, p < 0.001). Significant cortical areas encompassed the whole Superior Temporal Sulcus and Gyrus (STS/STG), bilaterally (Figure 5E). No significant results were found in V1 for the Low-frequency range (see Supplementary Figure 1; see Supplementary Figure 2 for results in the Temporal ROI).

### Envelope amplitude modulation across frequency ranges

Does the mapping of High and Low-frequency ranges differ? Within each ROI and hemisphere, we assessed whether the number of significant voxels associated with High and Low-frequency ranges, differed within each sound category and across all of them. Overall, the mapping of the two frequency ranges did not consistently vary in the occipital ROIs (see Supplementary Material). Conversely, in the bilateral Temporal ROI, the number of voxels encoding envelope variations in the High frequency range across all sound categories was significantly larger than the number of voxels explaining changes in envelope in the Low-frequency range (see Supplementary Material).

These results suggest that the functional organization of sound envelope mapping in V1 and Temporal cortex differs. Moreover, the fact that envelope PSD was greater in the Low compared to the High-frequency range for words, pseudowords, and artificial sounds (Figure 1B) further strengthen the evidence of a genuine sound envelope mapping in V1. Indeed, despite global signal regression, if sound envelope mapping in V1 would be the mere byproduct of arousal we should have observed a larger number of voxels mapping envelope PSD in the Low-frequency as compared to the High-frequency range (see Supplementary Material).

## Discussion

Here we investigated whether functional sensitivity to sound properties could be observed at the level of the striate cortex. To this aim, we measured the brain response to natural and synthetic auditory stimuli in the absence of visual input. In presence of a positive hemodynamic response, we found that the Calcarine cortex retained specific response patterns associated to our four sound categories. However, since a category-based organization could also be affected by multiple factors (e.g., imagery, semantics), we reconstructed envelope variations using activity of the Calcarine cortex (and Temporal ROI), thus ascribing its response to a fundamental acoustic feature. To this aim, we built an acoustic model based on the envelope power modulation in specific frequency ranges (Low, 2-6 and High, 6-10 Hz) and evaluated the cortical mapping of these features. In this context, categories were evaluated in isolation to investigate the degree of the response specificity and, because of their hierarchical organization, control for the contribution of multiple dimensions.

Amplitude modulations of sounds in both frequency ranges were traceable not only in temporal areas, typically involved in acoustic processing, but notably in the Calcarine cortex as well, indicating that sound amplitude modulations were spatially mapped in the earliest stages of the cortical visual pathway, and ultimately revealing a crossmodal representation of sound envelope.

The analysis of each sound category revealed that a crossmodal mapping in the striate cortex emerged in the absence of semantic content, as it was also found for meaningless speech stimuli, such as pseudowords. V1 involvement did not depend on the spectral properties of sounds: a representation of sound amplitude modulation emerged for artificial sounds in which the spectral properties had been flattened. Lastly, the tuning of this region was not speech-specific since a mapping appeared even for amplitude modulations of bird chirps. Overall, these results demonstrate that the human striate cortex represents sound envelope.

All the analyses performed in the current study are based on averaged representational spaces obtained from V1 activity through a group-level PCA, as well as averaged reconstructions of envelope properties by merging predicted models across individuals. Nonetheless, we also reported the overlap of single-subject within-category mapping of envelope in the Calcarine cortex (Supplementary Figures 5-8; from 0 to 20 individuals; p < 0.05, uncorrected for multiple comparisons). These findings further corroborate the group-level results, suggesting that envelope modulation of the striate cortex occurred in more than half of our participants.

### Semantic content and imagery

Evidence that sounds elicit specific functional activations in the primary visual cortex already exists. Muckli and colleagues (Muckli et al., 2015; Vetter et al., 2014) analyzed the hemodynamic signal of blindfolded individuals listening to complex auditory stimuli, such as the traffic noise or a forest auditory-scene. These authors showed that sound scene categories could be identified using decoding techniques from early visual cortex activity. Abstract conceptualization and visual imagery were suggested to represent the driving mechanisms of the visual cortex engagement; our results relative to isolated words may be interpreted similarly. However, a recent follow-up from the same group (Vetter et al., 2020) suggested that a similar successful decoding was also possible in early visual cortex of congenitally blind individuals, revealing that visual imagery was not necessary for the mapping of sound scenes in V1 and in other early visual areas. The present results of the multi-voxel analysis further support the possibility to dissociate in V1 specific patterns of category-based activity evoked by sounds.

Cross-modal recruitments have also been revealed in temporal regions using visual stimuli. The sight of scenes or written text carrying abstract acoustic features (e.g., a man playing the trumpet or the word “telephone”) were found to elicit responses in auditory regions (such as pSTG/MTG) as a function of the amount of acoustic relevance (Kiefer et al., 2008; Proverbio et al., 2011).

To control for the role of semantic information, we analyzed the mapping of pseudowords within our regions of interest. While in principle any auditory stimulus can elicit visual representations, these stimuli prevented coherent semantic-based imagery across participants. The observed responses to pseudowords provided evidence that semantic-related imageability alone could not explain the mapping of amplitude modulation of natural sounds observed in the striate cortex. Results provided by the artificial sound category further strengthened the lack of a role of semantics and imagery.

### Amplitude Modulation specificity

Brain areas such as STS are functionally tuned for the processing of speech-specific temporal structures. A recent study generated sound quilts by shuffling segments of a foreign language and other natural or artificial sounds to assess which regions were tuned for temporal properties (Overath et al., 2015). This approach allowed to preserve original sound features only at short timescales and disrupted them on longer timescales. While the primary auditory cortex was not sensitive to quilt durations, possibly due to the narrow frequency tuning which characterizes its response (Moerel et al., 2015; Thwaites et al., 2017), activity of the bilateral portion of STS varied as function of quilt lengths (Overath et al., 2015). Moreover, it was shown that portions of STG encode amplitude variations of speech and amplitude-modulated tone stimuli independent from other spectro-temporal features (Oganian and Chang, 2019). Taken together, these results reveal that regions coding speech-specific spectro-temporal features and non-speech-specific envelope variations coexist in the superior temporal cortex.

While our results could not exclude a role of spectral properties for the representation of sound features in the striate cortex, they reveal that natural, low-frequency modulations of the envelope power are sufficient to elicit selective responses in V1. Each artificial stimulus retained duration and estimated envelope of the word of origin, but its spectral structure was flattened. This approach ruled out the possibility that congruent variations of amplitude and of spectral properties explain the observed effects.

Finally, the sound envelope mapping observed in the striate cortex in response to artificial stimuli ensured that articulatory-based imagery does not explain our findings (Hauswald et al., 2018). While pseudowords are still reproducible from an articulatory point of view, artificial stimuli are not. Although each artificial sound was generated by a word, the flattening of spectral properties eliminated its intelligibility. Participants performed at chance level when asked to discriminate whether an artificial sound was derived by the original word or by the corresponding pseudoword (see Supplementary Figure 4B).

### The assessment of speech specificity

The human auditory system is functionally tuned for the processing of language since the early stages of development. Evidence exists that the auditory pathway is more tuned to language sounds than to environmental noise, even before it has reached full-term maturation (Webb et al., 2015). However, the degree to which temporal lobe regions are specific to speech processing remains to be ascertained. Evidence of speech-specific processing subregions of the superior temporal cortex was reported, mostly referring to the phonemic processing (Formisano, 2008; Mesgarani, 2014; Rampinini et al., 2019). STS tuning for spectro-temporal timescales appears to be selective for speech material. In case of natural, non-linguistic sounds were employed, such selectivity did not emerge (Overath et al., 2015). Nevertheless, examples suggesting that the temporal cortex is functionally tuned to sound features shared by both speech and non-speech materials are many. Tuning of STG for amplitude modulation of natural sounds was observed for both speech and non-speech sounds (Oganian and Chang, 2019).

To clarify whether the observed effects in V1 were functionally specialized for speech material, in our experiment non-linguistic natural sounds characterized by a rich amplitude-modulated profile, such as bird chirps, were presented as well. Intriguingly, V1 was recruited upon acoustic processing of this sound category as well, suggesting the lack of specificity for speech sounds. We ensured that none of our participants was expert in ornithology to avoid any imageability effect related to bird species. Indeed, while a generic visual representation of birds may have occurred in each participant, we observed a mapping of specific frequency ranges of the amplitude modulation.

### Lateralization

A left hemispheric functional specialization for speech processing exists already in three-month infants (Dehaene-Lambertz, 2002). Reviews of the literature have suggested that left hemispheric dominant responses progressively emerge in adults as acoustic materials increasingly provide speech-like input (Hickok, 2012; Peelle, 2012; Rauschecker and Scott, 2009). If amplitude-modulated noises activate bilaterally the primary auditory cortex, the processing of isolated phonemes and syllables typically results in activity along the left but not the right STS and MTG (DeWitt and Rauschecker, 2012). Moreover, the response tends to be greater in the left hemisphere as compared to the right when words and pseudowords are contrasted, suggesting a clear role of the left hemisphere for lexical processing (Davis and Gaskell, 2009). Furthermore, the hemispheric dominance in the superior temporal lobes seems to depend on the type of processing performed on speech signals. On the one hand, temporal details of speech signals primarily elicit brain activations of the left anterolateral STS and STG, whereas the right homolog is more sensitive to the spectral parameters (Obleser et al., 2008; Schönwiesner et al., 2005). Consistently, our results in the temporal cortex showed a major involvement of the left with respect to the right hemisphere, in both frequency ranges, only for word stimuli. The pattern of activity within the striate cortex did not reveal a consistent lateralization of the activity as a function of sound category with the exception of pseudowords, which for both frequency ranges, elicited a greater activity in the right than the left hemisphere.

### Temporal scale

Electrocorticography (ECoG) studies suggested a topographical representation of envelope power variations in STG and STS, with an anteroposterior gradient from Low to High modulation frequencies (Hullett et al., 2016), which depends on the spectral properties of sounds (Santoro et al., 2017). In the present study, due to the limited temporal resolution of the envelope variations associated to our stimuli, we could not assess whether the same gradient could be observed. However, a common pattern of activation in temporal areas emerged across all sound categories. A larger portion of the temporal cortex was associated with High compared with Low-frequency range, in both left and right hemispheres. Notably, in V1 the same type of organization did not emerge. When considering the mapping of all sound categories, no difference was found between the number of voxels associated with High and Low-frequency ranges, in either left or right V1. These results, together with the differences of laterality, suggest that V1 and the Temporal cortex do not share the same representation of sound amplitude variations.

### Functional significance in a multisensory frame of reference

Multisensory audio-visual stimulations subtly modulated the striate cortex activity in rodents (Ibrahim et al., 2016) and in awake monkeys (Wang et al., 2008). The anatomical scaffolding for early audio-visual interactions has been provided by tracing studies. Sparse monosynaptic (i.e., direct) anatomical projections originating from auditory areas and terminating in the primary visual cortex have been consistently found (Cappe and Barone, 2005; Charbonneau et al., 2012; Clavagnier et al., 2004; Falchier et al., 2002; Kim et al., 2015; Rockland and Ojima, 2003). In humans, the audio-visual convergence and integration between primary visual cortical areas were demonstrated at a functional level (Martuzzi et al., 2007; Mercier et al., 2013; Molholm et al., 2004; Romei et al., 2009). The existence of intrinsic functional coupling between primary visual and primary auditory areas was found even in the absence of external stimulation or task (Eckert et al., 2008). Modulations of hemodynamic activity measured in V1 have been associated to low-level audio-visual interactions (Watkins et al., 2006). Moreover, intracranial recordings revealed clear signs of audio-visual interplay in the striate cortex (Mercier et al., 2013). These consistent auditory modulations of visual cortical activity have provided a conceptual expansion of the range of potential activity patterns of the visual system. MVPA of fMRI signals first revealed that evoked responses of the primary visual cortex could be found even when auditory stimuli were applied transiently and in isolation. Distinguishable spatial patterns of neuronal responses could be elicited not only in the primary auditory cortex, but in V1 as well, showing that auditory inputs elicit a characteristic pattern of activation also in other primary sensory cortices (Liang et al., 2013). The observed patterns of crossmodal responses also include deactivations. Auditory looming stimuli have shown to induce widespread deactivations in primary visual cortex (Gau et al., 2020) that were maximal at the cortical surface and comprised regions that represent peripheral portions of the visual field. Importantly for the present study, these deactivations were found while participants attended to a fixation cross. Conversely, our results revealed an overall positive response in V1 for each sound category during auditory processing. The difference between these results might relate to the presence or absence of retinal input during sound processing.

Sound-driven crossmodal activations in the striate cortex have also been recently confirmed by measurements of intracranial stereotactic electroencephalographic (SEEG) recordings. High gamma neural oscillations were measured in striate cortex following the presentation of brief white noise stimuli. Such activity was found within the first 100 ms after stimulus onset and suggested the existence of populations of neurons performing auditory processing in striate cortex (Ferraro et al., 2020).

The crossmodal recruitment of V1 seems to particularly occur in case sight is missing. A large body of studies in blind individuals have reported V1 activations during sounds processing, including speech (see for reviews Pavani and Röder, 2012; Bedny et al., 2017; Fine et al., 2019; Ricciardi et al., 2020). It is noteworthy that early visual cortex of blind individuals synchronized to the envelope of intelligible speech, demonstrating to be associated to the processing of temporal dynamics of these signals (Van Ackeren et al., 2018). Moreover, short-term visual deprivation in sighted individuals is known to reveal activations in primary visual cortex during tactile exploration (e.g. Merabet et al., 2007). The present results indicate that in similar perceptual conditions, auditory tracking can be measured in sighted individuals as well, possibly suggesting that a rewiring following blindness might not be a prerequisite for the auditory-driven crossmodal recruitment to emerge.

Which neurophysiological mechanisms might explain the crossmodal modulation of V1 by an auditory input? Studies in the animal model have revealed that auditory stimulations change the firing and selectivity of V1 neurons. Specifically, a sound can modulate the responses to visual input by impacting inhibitory neurons in V1. In anesthetized mice, brief noise bursts (50 ms) generated activations in the auditory cortex, which in turn, had a modulatory role on inhibitory circuits in V1 and on the visually-driven spike activities (Iurilli et al., 2012). In particular, GABAergic inhibitory pyramidal neurons of the infragranular layers of V1 were activated by cortico-cortical connections originating in the auditory cortex (see Ibrahim et al., 2016).

One of the mechanisms which have been suggested as possible explanations for the crossmodal interactions in primary sensory cortices is the phase resetting of slow oscillatory activity (Kayser et al., 2008; Lakatos et al., 2008). Strong positive correlations exist between high frequency (40-130 Hz) local field potentials (LFP) and the fMRI signal. Also, negative correlations between low frequency (5-15 Hz) local field potentials (LFP) and the fMRI signal have been described (Mukamel et al., 2005). Both alpha and gamma oscillations are known to be linked to the activity of inhibitory GABAergic interneurons (see Jensen et al., 2012 for a review). The pattern in V1 observed in the present study could well be a byproduct of oscillatory activity occurring in the striate cortex. Whether such responses are mostly linked to slow, fast oscillations or their combination remains to be ascertained.

One possibility is that sound features, like the amplitude modulation, are conveyed to the visual cortex to prompt analyses on multisensory objects, as it occurs in the case of audio-visual associations, which are present already at the earliest developmental stages. At two months of age, infants show reliable matching of vowel information extracted in faces and voices (Patterson and Werker, 2003). It could be wondered whether the observed patterns of activation in V1 unveil common representations in auditory and visual cortices.

More than fifty years ago, Barlow (1961) suggested that sensory processing would exploit redundancies and the correlation structure existing within the input. Sensory signals are typically dynamic, multimodal streams. Correlation detection, such as lag and synchrony, has been suggested as a general mechanism behind multisensory integration (Parise et al., 2016). In speech, signal amplitude changes originate from the rhythmic movements of the mouth and the other phonatory apparatus (Plass et al., 2020). Importantly, coherent temporal modulations have been found between 2-7 Hz for both voice envelope and mouth openings (Chandrasekaran et al. 2009). The unity at the source level is reflected at the neural level. Speech signals conveyed by the sound envelope and rhythmic variations of lip movements are tracked by the neural oscillatory activity in both auditory and visual cortices (Giraud et al., 2012; Park et al., 2016). Moreover, auditory and visual tracking interact to maximize efficient processing (Crosse et al., 2016; Park et al., 2018). It is thus reasonable to speculate that the sound envelope mapping in V1 might represent input regularities which could contribute to efficient visual processing and ultimately to a unified audio-visual percept. In this respect, auditory amplitude modulated sounds have been reported to boost the visual-spatial frequency analysis, suggesting that a type of crossmodal mapping between these low-level auditory and visual features exists (Guzman-Martinez et al., 2012; Orchard-Mills et al., 2013).

It could also be wondered whether the observed patterns of activation in V1 unveil common representations between auditory and visual cortices. Amplitude modulations of natural sounds follow a 1/f relationship, which implies that low frequencies of amplitude modulation are the most represented. Noteworthy, also natural visual scenes respect the power law relationship of 1/f for the spatial and temporal luminance contrast (e.g., Srivastava et al., 2003). While the physical causes of the power law relationship in natural sounds and natural visual scenes occur for different reasons, it is possible that the two sensory systems create cross-modal (or modality-independent) feature maps (see for instance, Ricciardi et al., 2020; Heimler et al., 2020) of these types of fundamental stimuli properties already at the earliest stages of processing.

### Limitations

It could be wondered whether the associations between acoustic features and V1 activity could be explained by whole-brain hemodynamic fluctuations linked to arousal or alertness induced by sound onsets (Pisauro et al., 2016). It is worth mentioning that the global signal was responsible for up to 30% of the variance of our data. However, to exclude this confound the hemodynamic signal was corrected with a Global Signal Regression procedure (Aguirre, 1998; Macey et al., 2004). Thus, the observed pattern most likely represents auditory sensory information mapping in V1. As clearly shown in Figure 1B, the envelope PSD was greater for Low compared to the High-frequency range for words, pseudowords, and artificial sounds. If the sound envelope mapping in V1 would be the mere byproduct of arousal we would expect greater mapping in V1 for the Low compared to the High-frequency range. Results do not support this hypothesis, further strengthening the evidence of a genuine sound envelope mapping in V1. Nevertheless, further investigations are required to provide a finer characterization of the envelope PSD (e.g., with shorter frequency bins) associated to the Calcarine cortex hemodynamic activity.

In the present study, we focused on mapping of natural sound envelope and did not assess the role or spectral properties of sounds. Provided that high pitch and narrowband artificial sounds induce greater visual cortex excitability than lower pitch and broadband artificial sounds (Spierer et al., 2013), it could be hypothesized that spectral variations of natural sounds could be mapped as well in the striate cortex. Future studies might address this issue.

A further limitation of this study concerns the lack of evidence suggesting how sound envelope information reaches V1. It remains to be ascertained whether direct connections between auditory cortex and V1, or indirect routes involving relay regions, such as the parietal cortex, explain the observed patterns.

Finally, we selected relatively short (~1 sec) stimuli in isolation as they represented a compromise to estimate the elementary dynamic properties of natural sounds; in speech, typical syllabic and phonemic frequencies occur < 20 Hz. Ultimately, this allowed us to avoid the risks associated with the averaging of acoustic features of longer non-stationary sounds, which would be represented in the physiological hemodynamic response (de Heer et al., 2017). Moreover, this choice was made to prevent confounds associated with neural summations, which may potentially derive from the presentation of more continuous stimuli. However, further experiments are required to assess whether and to which extent the envelope of continuous stimuli would be mapped in V1 as well.

### Conclusions

The present study demonstrates that, in the absence of visual input, the human striate cortex maps sound envelopes. Ultimately, these findings strengthen the notion that not only multisensory processes occur in V1, but that even specific sound properties, such as amplitude modulation of natural sounds are represented within a brain area that once was conceived as purely visual.

## Materials & Methods

In the present study, we aimed at investigating the activity elicited by sounds in early visual cortex by using fMRI. We recorded four sets of sounds, three were speech-specific (i.e., words, pseudowords, and artificial stimuli, which were built upon the word category) and one included non-speech natural sounds (i.e., bird chirps; Figure 1). First, in the Calcarine cortex, we measured averaged BOLD (Blood oxygen level dependent) responses evoked by our sound categories as well as multivoxel patterns of brain activity through a group-level Principal Component Analysis (PCA; Figure 2). Then, from each sound wave, we calculated its envelope and the modulation power in the High (6-10 Hz) and Low (2-6 Hz) frequency ranges. Hence, within each sound category, we exploited voxel-wise modeling to measure the association between these acoustic features and the hemodynamic activity in occipital and temporal areas (Figure 3).

### Subjects

Twenty right-handed healthy individuals were recruited for the fMRI study (10F; mean age ± standard deviation: 34.5 ± 6.5 years; Edinburgh Handedness Inventory: 16.4 ± 2.5 (Oldfield, 1971). All participants were native Italian speakers, recruited from the local area of Lucca. None of the participants had been diagnosed with language-related developmental disorders (e.g., dyslexia, specific language impairment, delay of language onset). None of the subjects had any history of any relevant medical, neurological, or psychiatric condition that could affect brain development or function. None of them was taking any medication at the time of the study. Prior to the enrollment to the study, they gave their written informed consent and received 50 euros at the end of the experiment as a reimbursement. Ethical approval was obtained from Area Vasta Nord Ovest Ethics Committee (Protocol number 1485/2017) and conducted according to the Declaration of Helsinki (2013).

### Stimuli

First, we selected 20 tri- and quadri-syllabic commonly used Italian words belonging to the same semantic category -i.e., vegetables; word length: 8 ± 2; CoLFIS frequency: 1.25 ± 1.74 (Bambini and Trevisan, 2012)-. Words pertained to graspable objects with comparable sizes. Starting from these 20 words, we created 20 pseudowords using “Wuggy”, a multilingual pseudoword generator able to produce polysyllabic stimuli that respect the phonotactic constraints of the Italian language (Keuleers and Brysbaert, 2010). All the pseudowords had no meaning and matched the corresponding word stimuli in the sub-syllabic structure, the letter length, and the length of subsyllabic segments, besides matching transition frequencies of the reference word (see Supplementary Table 1). Our speech-specific stimuli were read by a trained Italian actress and were acquired in a recording studio, using a microphone (Behringer C-1U; 40-20,000 Hz, 130db max SPL) connected to an iMac™ (Apple Inc.). The sampling frequency of the recording was 44100 Hz. The resulting waveforms were edited using the Audacity software (©Audacity Team) to remove silence periods before and after each stimulus, and were slightly changed in tempo (within ±15%) to set the duration at 1 s for all stimuli. As result, sounds lasted about 1.00 ± 0.06 s for words and 1.03 ± 0.06 s for pseudowords. Afterward, we manually tagged the duration of syllables and phonemes composing our words and pseudowords (medians and confidence intervals estimated by bootstrapping method, 1000 iterations).

Then, we constructed a third class of stimuli (i.e., artificial sounds), reproducing in detail the low-level features (duration and estimated envelope) of the above-mentioned words, but with a completely different spectral structure. As detailed below, we aimed at creating a set of stimuli that retained the prosodic pitch modulation of the original words, but, at the same time, fine spectral details were flattened (Figure 1A). First of all, the original audio trace of each word was decomposed into 30 critical bands linearly spaced in the Equivalent Rectangular Bandwidth scale between 100 and 8000 Hz, using a zero-phase gammatone filter-bank (Hohmann, 2002). Single sub-band envelopes were then evaluated as the amplitude of the corresponding Hilbert transform and linearly summed across critical bands. After being smoothed with a 10ms-moving average filter, the obtained global envelope was used to modulate the amplitude of a spectrally-homogeneous source. This source was generated by filtering white Gaussian noise in a single critical band of the gammatone filter-bank whose central frequency was characterized by the lowest absolute threshold of hearing (f = 3.434 kHz; see Figure 1A). The absolute threshold of hearing within the 30 central frequencies was evaluated considering a diffuse binaural field, according to the ISO 389-7: 2005 standard. All the described operations were performed within the Auditory Modeling Toolbox (Søndergaard and Majdak, 2013) in the Matlab environment.

As an additional control condition, we selected bird chirps. They represent naturalistic sounds, comprising envelope modulations in the low-frequency bands (<20 Hz) resembling human speech. In particular, we extracted 20 bird chirps of 10 different bird species from video documentaries retrieved from YouTube® (128 kbps, compressed using advanced audio coding and converted to 44100 Hz) (see Supplementary Table 1). Bird chirps matched the duration of the other sound categories (1.01 +0.02 s).

Loudness was normalized across all stimuli by imposing a fixed root mean square value on each raw signal.

### Feature extraction: modulation power in the Low and High-frequency ranges

In order to describe each stimulus, two separate features were extracted from the signal envelope, conveying information about the Low (2-6 Hz) and High (6-10 Hz) frequency ranges of the modulation power, which were associated with phonemic and syllabic phonological properties in the speech material. In particular, the signal envelope was calculated using the procedure described above to generate artificial stimuli (Biesmans et al., 2015; Hohmann, 2002). Syllabic and phonemic power was then defined as the integral of the envelope power spectral density (PSD) in the two ranges of interest. These two intervals were centered on the manually estimated syllabic and phonemic frequencies, reported in the Results section. For consistency, the amplitude variations in the same Low and High-frequency ranges were also extracted for bird chirps. Importantly, the PSD in Low and High frequency ranges were uncorrelated within each sound category (word: Pearson’s r=0.213, CI-95^th^: −0.128 - 0.534; pseudoword: r=0.257, CI-95^th^: −0.294 - 0.616; artificial: r=- 0.025, CI-95^th^: - 0.359 - 0.342; bird chirps: r= −0.222, CI-95^th^: −0.523 - 0.175).

Following the stimuli generation procedure words and pseudowords had comparable envelope power modulations in the Low-frequency range (Pearson’s r=0.375, CI-95^th^: 0.061-0.640), whereas they differed in the High-frequency range (r=0.189, CI-95^th^: −0.259-0.543). This was due to the preservation of the syllabic rate and to the alteration of the phonemic features. Conversely, since words and artificial sounds shared the same envelope, the two ranges of the envelope PSD were highly collinear (2-6 Hz range: Pearson’s r=0.963, CI-95^th^: 0.909-0.986; 6-10 Hz range: r=0.886, CI-95^th^: 0.721-0.982).

### Experimental procedures

A slow event-related paradigm was implemented using E-prime 3.0 software (Psychology Software Tools, Sharpsburg, Pennsylvania), and comprised 80 stimuli (equally divided across the four categories, i.e., 20 words, 20 pseudowords, 20 artificial sounds, and 20 bird chirps). Each event comprised a ~1 s stimulus followed by 9 s of rest. Each stimulus was randomly presented three times across six runs lasting 8 minutes each (Mumford et al., 2014). Participants laid down blindfolded in the scanner. To ensure participant’s attention, they were instructed to detect by a button press rare target sound. These rare deviant sounds (30 out of 270) were generated adding to our original stimuli a silent period (lasting 550 ms, starting at 150 ms after stimulus onset), and were distributed in time in a pseudo-random order to ensure the presence of ten targets across two sequential runs. The aim of the task was also to ensure that each participant maintained the attentional focus on the temporal dynamics of sound waves. Behavioral responses were analyzed to calculate precision -i.e., True Positive: TP; False Positive: FP; False Negative: FN; TP/(TP+FP)- and recall –i.e., TP/(TP+FN)- measures. The behavioral task resulted in a high level of precision (87% ± 3%) and recall (90% ± 3%), suggesting that participants attended to the stimuli. During the scanning session, prior to the fMRI acquisition, participants underwent a brief session (~12 minutes) to familiarize themselves with the task.

#### MRI Data Acquisition and Preprocessing

Brain activity was recorded using a Philips 3T Ingenia scanner, equipped with a 12 channels phased-array coil, and a gradient recall echo-planar (GRE-EPI) sequence with the following acquisition parameters: TR/TE = 2000/30ms, FA = 75°, FOV = 256 mm, acquisition matrix = 84 × 82, reconstruction matrix = 128 × 128, acquisition voxel size = 3 × 3 × 3 mm, reconstruction voxel size = 2 × 2 × 3 mm, 38 interleaved axial slices, 240 volumes. Twenty seconds of rest preceded in each run the first stimulus onset and followed the last one. Stimuli were delivered with MR-compatible on-ear headphones (VisuaStim, 30 dB noise-attenuation, 40 Hz to 40 kHz frequency response).

Three-dimensional high-resolution anatomical image of the brain was also acquired using a magnetization-prepared rapid gradient echo (MPRAGE) sequence (TR/TE = 7/3.2ms, FA = 9°, FOV = 224 mm, acquisition matrix = 224 × 224, voxel size = 1 × 1 × 1 mm, 156 sagittal slices).

The fMRI preprocessing was performed with the AFNI software package (Cox, 1996). All volumes within each run first underwent spike removal (*3dDespike*), were temporally aligned (*3dTshift*), and corrected for head motion (*3dvolreg*). The transformation matrices were also used to compute the frame-wise displacement (Power et al., 2012) that identified time points affected by excessive motion (threshold: 0.5 mm). Afterward, spatial smoothing was performed with a Gaussian kernel having 4 mm Full Width at Half Maximum (FWHM). In this regard, we adopted the AFNI’s *3dBlurToFWHM* tool, which first estimated and then iteratively increased the smoothness of data until a specific FWHM level was reached. Considering that the original smoothness was above 3 mm (*3dFWHMx*), this procedure generated a final homogeneous smoothness of 4 mm across voxels and subjects, far less than the one obtained by simply adding a smoothing filter of the same width to the data and aimed at preserving the functional distinctions across adjacent sulcal walls and sulci. Runs were normalized by dividing the intensity of each voxel for its mean over the time series. Normalized runs were then concatenated and a multiple regression analysis was performed (*3dDeconvolve*). Each stimulus comprised three repetitions and was modeled by a standard hemodynamic function (i.e., BLOCK), lasting 1 s. Deviant sounds and button presses were similarly modeled but were excluded from the multivariate procedure detailed below. Movement parameters, frame-wise displacement, and polynomial signal trends were included in the analysis as regressors of no interest. The t-score maps of the 80 stimuli were used as input in the multivariate analysis. Single-subject data were also registered to the MNI152 standard space (Fonov et al., 2009) using a nonlinear registration (AFNI *3dQWarp*) and resampled to a final resolution of 2×2×2 mm.

#### Regions of interest

We focused our analysis on two regions of interest in temporal and occipital cortical areas. For temporal areas, we adopted the AICHA atlas (Joliot et al., 2015), which takes into account brain hemispheric specialization and is widely used to identify language responsive areas. The selected regions were the bilateral Superior Temporal Gyri and Sulci as well as the Middle Temporal Gyri. These regions were selected to isolate the portions of the auditory stream activated during the processing of fine spectral features (Santoro et al., 2017), from short-timescales (e.g., phonemes) up to more complex sound patterns (e.g., syllables and words) (DeWitt and Rauschecker, 2012; Keitel et al., 2018; Pernet et al., 2015). Since the primary aim of our study was to investigate the activity elicited by sounds in the early visual cortex, we selected the Calcarine cortex as a region of interest, using the probabilistic map (V1, threshold > 10%; Gau et al., 2020) by Wang and colleagues (Wang et al., 2015). A spatial mask was applied to temporal and occipital areas to select gray matter voxels only (gray matter probabilistic threshold > 0.25).

#### Pattern analysis in the Calcarine cortex

We first measured the overall BOLD activity in bilateral V1, averaging at the single participant level, across stimuli of each sound category. Sound evoked activity was compared to baseline using a one-sample two-tailed Student’s t-test (dofs = 19). Results were reported in Figure 2A. Then, we analyzed the response patterns of V1 by means of a group-level PCA. First, in the bilateral Occipital ROI, we performed a PCA in each participant by decomposing voxels activity in a fixed set of Principal Components (PCs). The final number of PCs (n = 63) was selected to retain 95% of cumulative explained variance averaged across individuals. In each participant, all the retained PCs were z-scored. Subsequently, the PCs of all individuals were aligned using Procrustean transformation in a two-pass procedure (Haxby et al. 2011; Lettieri et al., 2019). In the first pass, the PCs from all subjects were compared to each other using Procrustean transformation. This procedure generated a set of sums of squared errors (i.e., goodness-of-fit) for each participant pair. Hence, we identified the reference individual by selecting the participant with the lowest sum of squared errors across pairs (i.e., centroid individual). Then, in the second pass, we computed again Procrustean transformations to align each individual’s PCs to the corresponding PCs of the reference participant. Then, PCs were averaged across individuals to obtain a group-level hyper-aligned common space (Haxby et al. 2011). Subsequently, from each hyper-aligned PC a Representational Space (RS) was obtained by using the Euclidean distance as measure of pattern similarity. Finally, we evaluated in each RS, the Michelson’s coefficient as a measure of contrast for each category (Michelson, 1927):

where D_between_ and D_within_ represented the mean between- and within-category distance, respectively. A contrast of zero indicated that a category-based representation was not present in the RS, whereas a positive measure (up to 1), indicated that the stimuli pertaining to a category of interest retained a lower distance (i.e., high similarity) compared to the distance between stimuli of the other categories. Therefore, we obtained a measure of category contrast from each RS and its related PC. To test the significance of that estimate, we performed a permutation test, by shuffling rows and columns of the participants PCs and by repeating the entire procedure (i.e., identification of the reference individual, alignment of PCs of each participant to the reference and the evaluation of the contrast) 10,000 times, thus obtaining a null distribution of coefficients for each RS and category which was compared to the real contrast value. Results were corrected for multiple comparisons using False Discovery Rate (FDR) to control for test dependency (Figure 2B, Benjamini and Yekutieli, 2001). For visualization purposes, we also averaged the RS generated from each significant PC (Figure 2C) to highlight a category-based representation in the Calcarine cortex. However, a category-based organization could result from multiple factors (e.g., imagery, semantics, different level of arousal associated to each category) rather than being a genuine representation of acoustic features. Therefore, in the subsequent analyses we adopted a within-category approach, aiming at reconstructing envelope variations from brain activity, and conducted the analysis at voxel level to provide a finer spatial description of sound envelope mapping.

#### Global activity regression

To exclude that brain-wide hemodynamic fluctuations could impact the results obtained in temporal and occipital cortices, we opted for a global signal regression procedure (Aguirre, 1998; Macey et al., 2004). The rationale behind this approach is related to the fact that global hemodynamic activity represents motion, vascular, cardiac, and respiratory confounds, as well as sources of neural activity (Liu et al., 2017). Intriguingly, the latter components were described as task-related anticipatory mechanisms (Cardoso et al., 2012; Sirotin and Das, 2009), variations in arousal or alertness (Pisauro et al., 2016), and produced large-scale hemodynamic responses also in resting state, particularly in the eyes-closed condition (Scholvinck et al., 2010). Thus, to avoid possible effects of these physiological confounds, we removed the global signal from our data. The global activity of the full set of stimuli was obtained by averaging hemodynamic responses across gray and white matter voxels. Prior to the regression procedure, we first assessed whether the four categories significantly retained a different whole-brain average activity (see Supplementary Figure 3A). The results of this procedure did not highlight specific category-based differences, suggesting that global activity retained a common, positive, hemodynamic average response. Second, we measured the correlation (Spearman’s ρ) of the global signal between each pair of subjects (see Supplementary Figure 3B). The results demonstrated that global activity had a subject-specific pattern (ρ=0.002, CI-99^th^: −0.258, 0.292). We directly measured the association between global activity and envelope modulation power in the High and Low-frequency ranges using Spearman’s ρ across participants (see Supplementary Figure 3C). The results of this analysis demonstrated that global activity was, on average, uncorrelated with the modulation power of our stimuli. To rule out a possible correlation at the single-subject level, global responses were regressed out across all voxels (Macey et al., 2004). The association between global and single-voxel activity, measured using R^2^, was reported in Supplementary Figure 3D, and showed a relatively high impact of the global signal removal in primary sensory regions, ranging on average between R^2^≈0.2 to R^2^≈0.3 in the occipital ROI.

#### Reconstruction of envelope features from brain activity

After the removal of the global brain signal, the obtained hemodynamic responses were associated with the envelope modulation power using a searchlight approach (Kriegeskorte, 2006) and a machine learning algorithm based on principal component regression (Thirion, 2014). Specifically, for each stimulus category, a searchlight analysis was performed in the above-mentioned regions of interest to predict envelope features using principal components derived from brain activity (see Figure 2A).

It is important to note that, despite the analysis was performed in volumetric space, we respected the cortical folding by sampling voxels (radius: 8 mm) along the cortical ribbon (i.e., the space between pial surface and gray-to-white matter boundary) (Yu, 2019). Specifically, we ran the Freesurfer (version 6.0.0) recon-all analysis pipeline (Reuter et al., 2012) on the standard space template (Fonov et al., 2009), used as a reference for the nonlinear alignment of single-subject data. This procedure provided a reconstruction of the gray-matter ribbon, which has been used to isolate searchlight voxels taking into account the cortical folding. Voxel proximity was evaluated using the Dijkstra metric as it represents a computationally efficient method to estimate cortical distance (Fischl, 1999; Lettieri et al., 2019).

Similarly to the procedure adopted for the multivariate pattern analysis, for each searchlight, we performed a PCA to extract orthogonal dimensions from normalized voxel responses within each experimental condition and subject. The retained PCs explained on average 95% of the total variance across the regions of interest and subjects. Afterward, we performed a multiple regression analysis, using the PC scores as the independent variable and the power of Low (e.g., 2–6 Hz) and High (e.g., 6-10 Hz) modulation frequencies as the dependent one. To avoid overfitting and to obtain a robust estimate of these associations (Varoquaux et al., 2017), we performed the regression using a 10-fold cross-validation procedure on both envelope frequency ranges separately. This procedure, ultimately led to a reconstructed model of predicted power values in each experimental condition, subject and searchlight. The reconstructed models were compared to the actual ones by calculating their root mean squared error (RMSE; Poldrack et al., 2019).

To measure the statistical significance, w**e** first averaged the reconstructed models across participants in each experimental condition and searchlight, thus obtaining a group-level predicted set of acoustic features. Secondly, using the same procedure of the multivariate pattern analysis, we performed a non-parametric test (1,000 iterations), by shuffling the independent variable (i.e., PC scores) in the training set of each k-fold iteration, keeping the permutation scheme fixed across voxels (Winkler, 2016). This procedure led to a set of 1,000 group-level reconstructed models for each experimental condition and searchlight and ultimately provided a null distribution of RMSE coefficients, against which the actual association was tested. To compute a more accurate estimate (i.e., beyond the number of iterations used in the non-parametric test) of the p-value, we modeled the left tail of the RMSE null-distribution (p < 0.10) using a generalized Pareto distribution (Knijnenburg et al., 2009; Winkler, 2016). Results were corrected for multiple comparisons using FDR for each hemisphere separately to control for test dependency (Benjamini and Yekutieli, 2001).

Finally, we measured through a series of χ^2^ tests, the difference between volumes comprising significant voxels across Low and High-frequency ranges, sound categories and ROIs. Since the number of voxels in the ROIs exceeded the real number of degrees of freedom (dofs), we estimated the latter by means of a Monte Carlo simulation. Specifically, since our results were obtained using both a voxel level statistical threshold and a minimum cluster size of 20 voxels, we repeated random sampling of patches of contiguous cortex with a volume of 20 voxels until the entire region of interest was covered. This procedure was repeated 1,000 times. Then, we considered the average number of iterations necessary to fill the region of interest as a good approximation of its dofs. All the analyses were performed using MATLAB R2016b (MathWorks Inc., Natick, MA, USA).

## Data availability

The data that support the findings of this study will be provided to all readers upon reasonable request.

## Code availability

All relevant MATLAB code is available from the corresponding author upon reasonable request.

## Acknowledgements

We thank Russel Poldrack for suggestions concerning data analysis. We also thank Davide Crepaldi for proving access to the Italian version of the Wuggy software.

D.B. was funded by PRIN 2017 research grant. Prot. 20177894ZH.

## Author contributions

A.M., G.H, A.L. and D.B. conducted the experiments; A.M., G.H, M.B., and D.B. designed the experiments and wrote the paper. All authors revised the manuscript.

