## Supplementary Material for "Auditory features modelling reveals sound envelope representation in striate cortex"

### Supplementary Figures

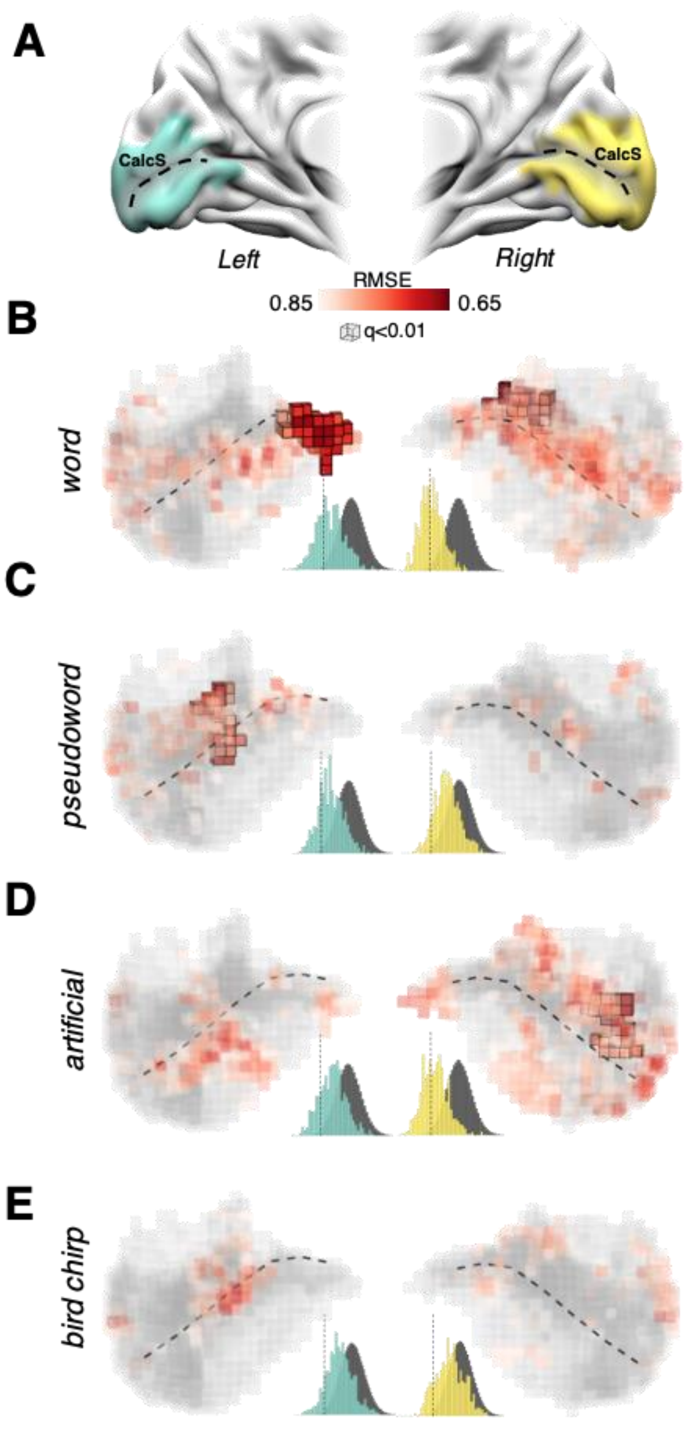


**Supplementary Figure 1. Reconstruction of the sound envelope power in the 2-6 frequency range in V1.** (A) V1 ROI in the left (cyan) and right (yellow) hemispheres. (B, C, D, E) Reconstruction in V1 of the sound envelope power in the Low (2-6 Hz) frequency range as a function of sound categories (i.e. word, pseudoword, artificial and bird chirps). Significant voxels surviving the multiple comparisons correction are represented by highlighted cubes (q<0.01; minimum cluster size = 20 voxels), whereas colored cubes displayed uncorrected threshold results at p<0.01. Population histograms represent the overall reconstruction performance in the left (i.e., cyan histogram) and right (i.e., yellow histogram) hemispheres for each ROI and sound category as compared to their null distributions (i.e., dark grey histogram; dashed vertical lines in histograms represents the $1^{st}$ percentile). Results show that V1 represents sound envelope power for all sound categories with a lower RMSE as compared to the null distribution. See Supplementary Figures 6 and 8 for single subjects results. For anatomical landmarks please refer to Figure 2. (B) For the word category, an involvement of the Calcarine cortex was found in the Low frequency range (with no difference between overall engagements of left and right hemispheres; left ~ 3.3% and right ~ 2.2% of the total volume; number of voxels in left vs. right hemisphere: $\chi 2$ (1, 334) = 0.475, p = 0.49).

(C) As for the High-frequency range, the comparison between the number of voxels of V1 associated with the envelope power of pseudowords in the Low-frequency range was greater in the left compared to the right hemisphere (left ~ 2.1% and right = 0% of the total volume; $\chi 2$ (1, 334) = 4.169, p = 0.041).

(D) Consistently with the High-frequency range, no difference emerged between left and right hemispheres involvement (left = 0% and right ~ 1.8% of the total volume; $\chi 2$ (1, 334) = 2.938, p = 0.086), confirming that the modulation of V1 activity associated to the artificial sounds was not selective for the 6-10 Hz frequency range. (E) No significant results were found in V1 for the Low-frequency range in the Bird chip category.

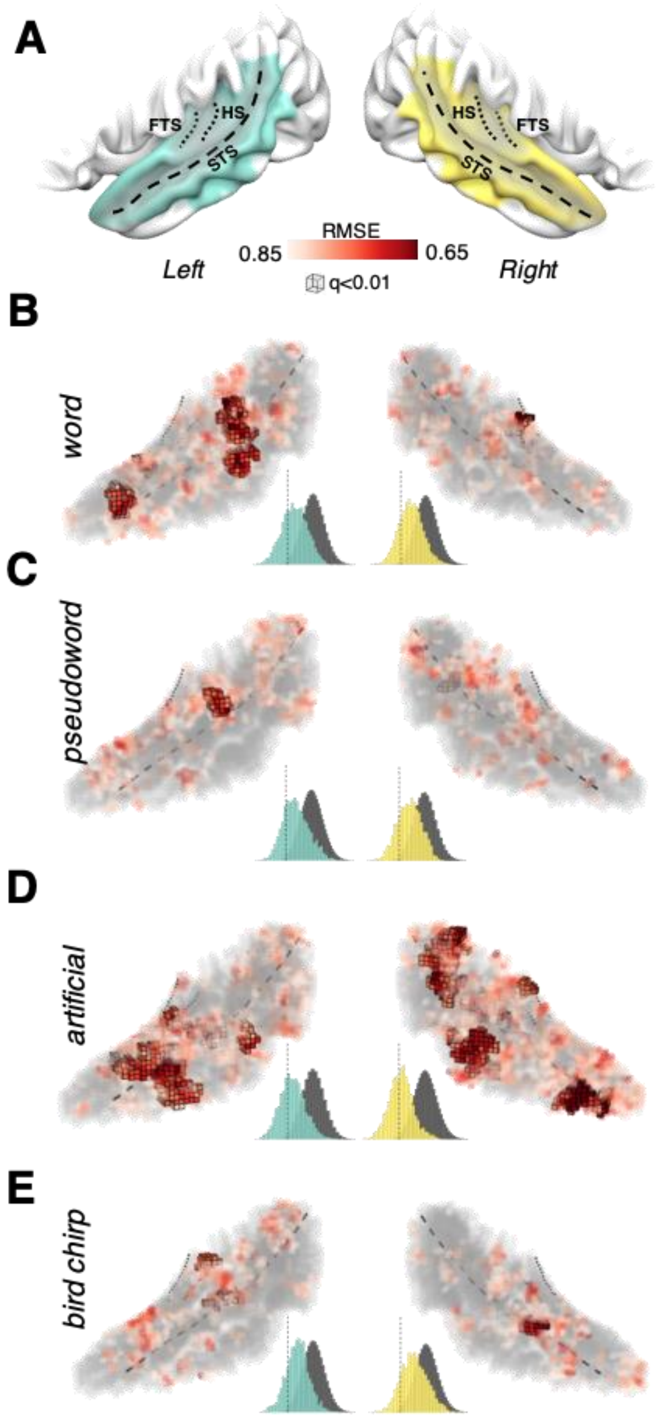


**Supplementary Figure 2. Reconstruction of the sound envelope power in the 2-6 Hz frequency range in the Temporal ROI.** (A) Temporal ROI in the left (cyan) and right (yellow) hemispheres. (B, C, D, E) Reconstruction in the temporal ROI of the sound envelope power in the Low (2-6 Hz) frequency range as a function of sound categories (i.e. word, pseudoword, artificial and bird chirps). Significant voxels surviving the multiple testing correction ($q<0.01$; minimum cluster size = 20 voxels) are represented by highlighted cubes, whereas colored cubes displayed uncorrected threshold results at $p<0.01$. Population histograms represent the overall reconstruction performance for each ROI and sound category as compared to null distributions. For anatomical landmarks please refer to Figure 2. (B) For the word category, an asymmetry in the temporal cortical areas were found (left ~ 4.4% and right ~ 0.8% of the total volume; $\chi 2$ (1, 1615) = 22, $p < 0.001$). (C) Significant patches of cortex were identified in the bilateral mSTS/mSTG (see Supplementary Figure 2) with no hemispheric dominance for the pseudoword category (left ~ 0.4% and right ~ 0.3% of the total volume; $\chi 2_{(1, 1615)} = 0.200$, $p = 0.654$). (D) Greater involvement of the right hemisphere compared to the left hemisphere was found for the artificial category (left ~ 5.1% and right ~ 7.5% of the total volume; $\chi 2$ (1, 1615) = 4.186, $p = 0.041$) Significant patches of temporal cortex were bilaterally identified in the anterior part of the Superior Temporal Sulcus and Gyrus (aSTS/aSTG), as well as the left mMTG and in the right mid-portion of STG (mSTG) and pSTS (see Supplementary Figure 2). (E) Conversely, for the bird chirp category, small clusters of voxels in the temporal cortex were associated with envelope characteristics in the Low-frequency range: ~1% of the voxels in the left hemisphere and ~0.4% in the right hemisphere, with no hemispheric dominance ($\chi 2$ (1, 1615) =2.288, $p = 0.13$). Significant clusters encompassed the middle Superior Temporal Sulcus and Gyrus (mSTS/mSTG), bilaterally.

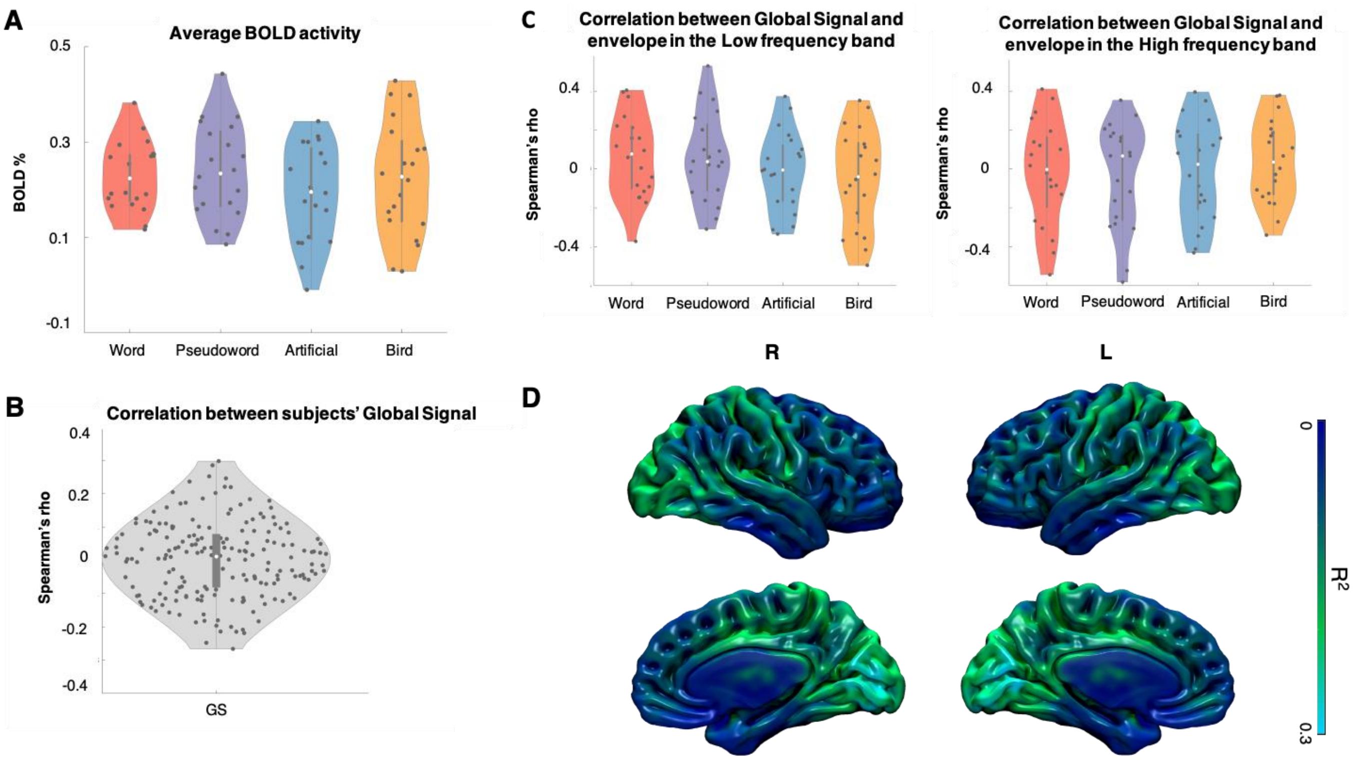


**Supplementary Figure 3. Average BOLD activity for each stimulus and Global Signal Regression.** (A) Average BOLD activity in grey and white matter voxels for the four experimental conditions is reported. Each dot represents a stimulus. Specifically, BOLD % signal change was (mean±std, min-max) 0.23±0.07, 0.12-0.38 for word category, 0.24±0.10, 0.09-0.44 for pseudoword category, 0.19±0.10, -0.01-0.34 for artificial category, 0.22±0.12, 0.03-0.43 for bird category. Paired t-test demonstrated no significant differences in baseline activity between the four categories (dof=19, all comparisons p>0.05). (B) Correlation of the average BOLD activity in grey and white matter voxels between each pair of subjects considering all the 80 stimuli. Results showed that global BOLD activity was not correlated among subjects with a Spearman's σ (mean±std, min-max) 0.00±0.11, -0.27-0.30. T-test further supported that average group correlation was not different from zero (dof=189, p=0.822). (C) Association between brain average BOLD activity and the modulation power in the low frequency range (i.e., 2-6 Hz for word, pseudoword, artificial categories and 4-8 Hz for the bird one) and in the high frequency range (i.e., 6-10 Hz for word, pseudoword, artificial categories and 8-12 Hz for the bird one) as well. Each dot represents a stimulus. Results indicated that overall our feature sets were not correlated with global brain activity. Specifically, for the low frequency range, word category retained a Spearman's ρ (mean±std, min-max) 0.06±0.21, -0.37-0.40, pseudoword 0.06±0.22, -0.31-0.53, artificial -0.01±0.20, -0.34-0.37 and bird -0.05±0.25, -0.49-0.35. Moreover, for the high frequency range, word category retained a Spearman's ρ (mean±std, min-max) -0.02±0.27, -0.54-0.41, pseudoword -0.03±0.27, -0.58-0.35, artificial -0.01±0.26, -0.43-0.39 and bird 0.04±0.21, -0.34-0.38. T-tests further supported that these associations were not different from zero (dof=19, all tests p>0.05). (D) Global signal regression procedure, as describe in the Methods section of the main manuscript. Moreover, we mapped the $R^2$ coefficient to measure the impact of global activity on single voxels. Results showed a relatively high impact of this procedure, ranging on average between $R^2 \approx 0.2$ to $R^2 \approx 0.3$ in the occipital and parietal lobes.

Overall these supplementary results showed the effects of global signal in primary sensory regions, demonstrating at the same time that 1) global activity was not associated to different BOLD responses across the four categories, 2) it did not retained a specific pattern of activity across subjects, and 3) it was not even correlated to our sound descriptors. For these reasons it was unlikely that the association between envelope features and BOLD activity in early visual cortex may be driven by spurious global confounds (e.g., motion, vascular components) or sources of large-scale neural activity (e.g., arousal, alertness).

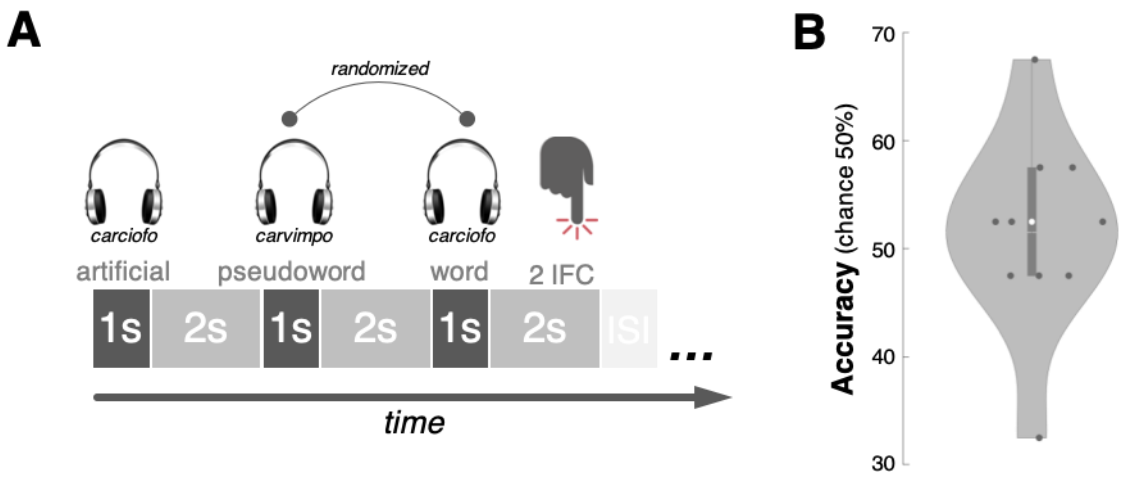


**Supplementary Figure 4. Testing the intelligibility of artificial sounds category.** (A) To test the intelligibility of the artificial stimuli, participants listened to each artificial sound and were then asked to identify the original sound from which it was generated by means of a 2-Alternative Forced-choice task. For an artificial sound generated starting from a word (e.g. the artificial sound derived from "*carciofo*"), we asked to identify the original sound choosing between the word ("*carciofo*") and its associated pseudoword ("*carvimpo*"). (B) The results demonstrated that accuracy was at chance level (accuracy ± SE: 51.5% ± 3%, t(9)=0.52, p=0.3069, one-tail t-test), providing evidence that intelligibility was not retained despite the envelope modulations were maintained.

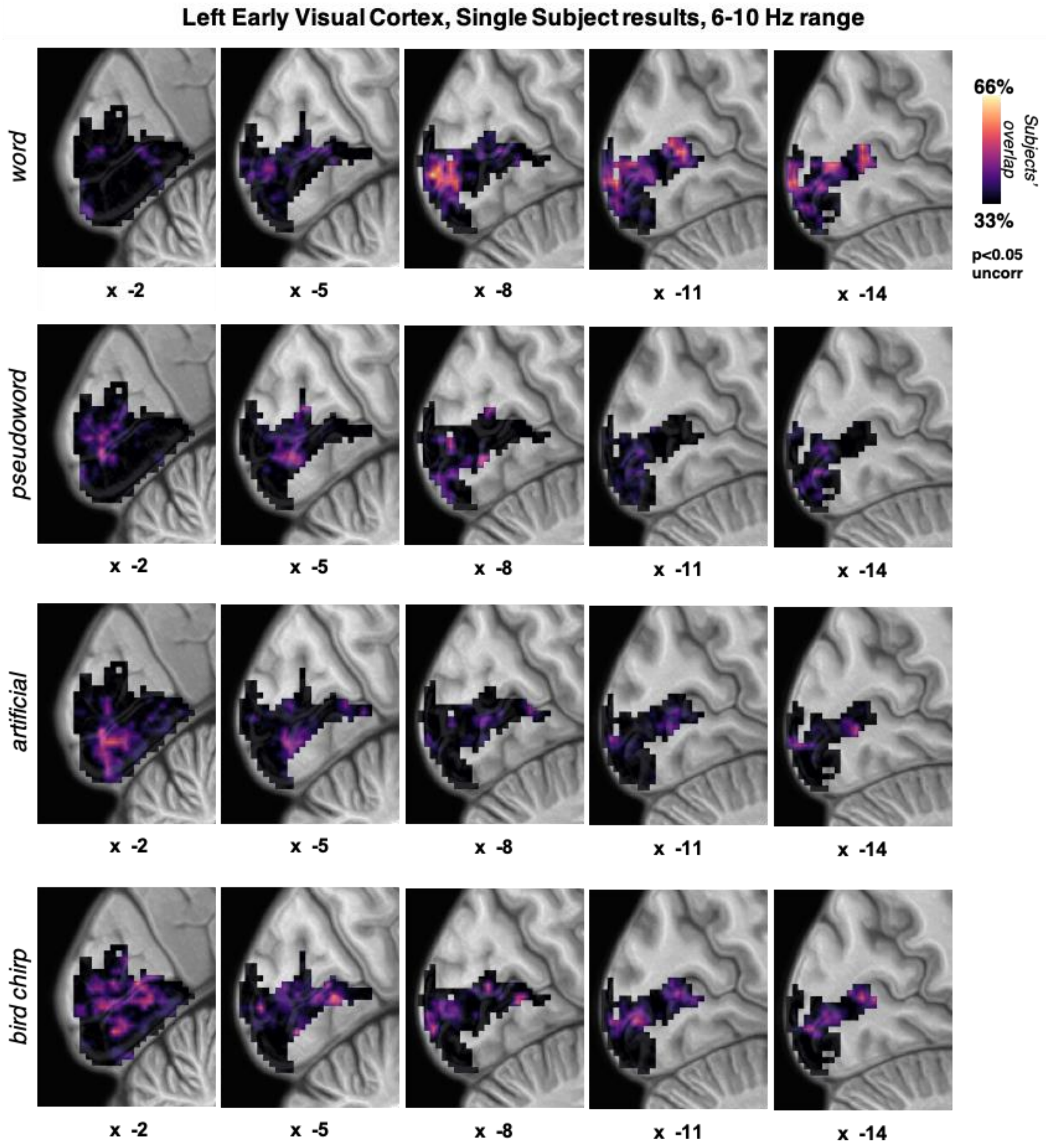


**Supplementary Figure 5. Envelope mapping single-subject results in the 6-10 Hz frequency range in left early visual cortex.** Within-category results of the single-subject envelope mapping in left Calcarine cortex are reported, representing the spatial overlap between participants (from 0 to 20 individuals; $p < 0.05$, uncorrected for multiple comparisons). Darker colors represent less overlap (starting from 33% of overlap between participants), brighter colors represent for up to 66% of overlap between participants.

Each row shows envelope mapping within each category (i.e. word, pseudoword, artificial and bird chirps), moving along the x-axis (from medial slices to lateral ones).

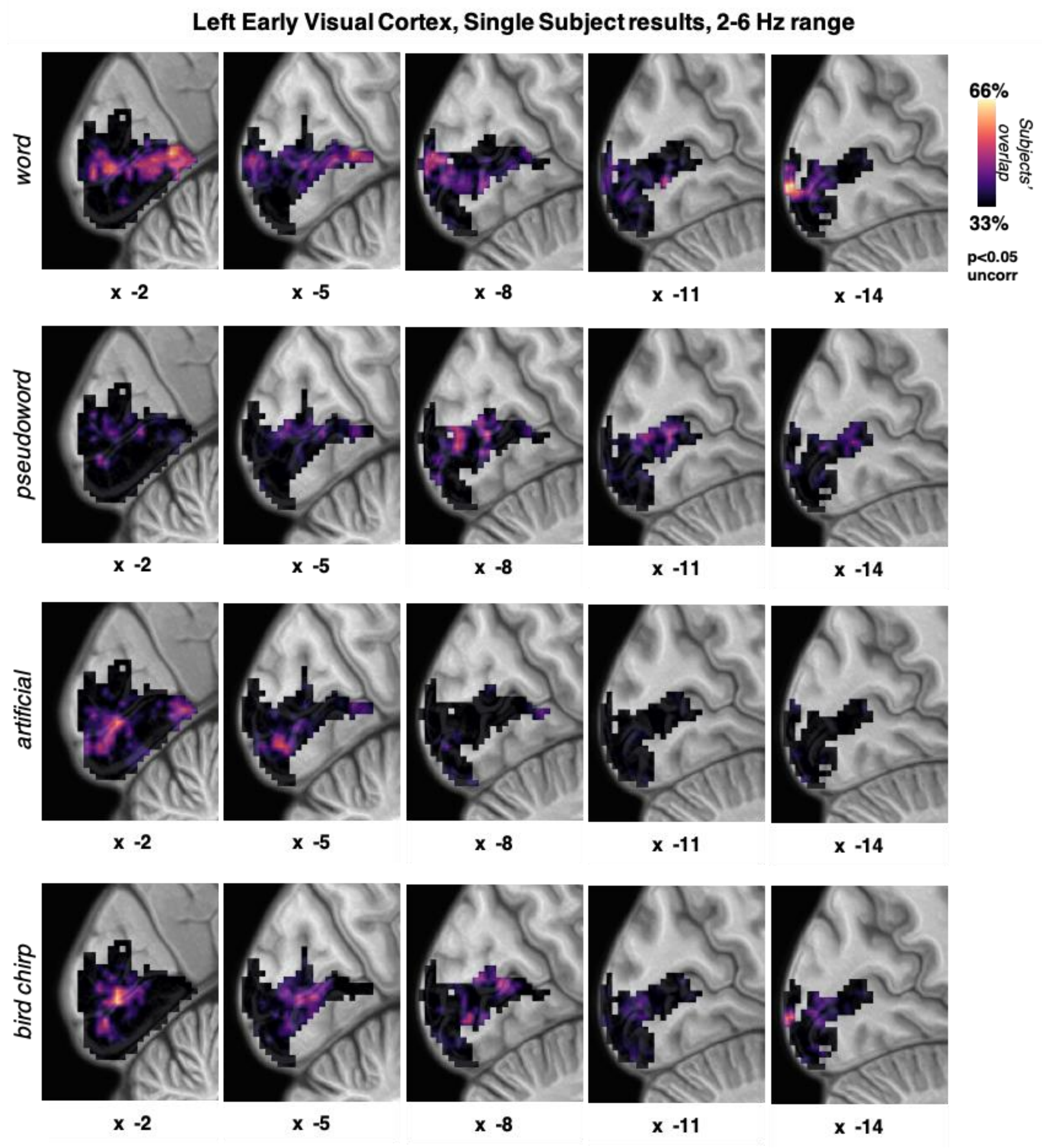


**Supplementary Figure 6. Envelope mapping single-subject results in the 2-6 Hz frequency range in left early visual cortex.** Within-category results of the single-subject envelope mapping in left Calcarine cortex are reported, representing the spatial overlap between participants (from 0 to 20 individuals; $p < 0.05$, uncorrected for multiple comparisons). Darker colors represent less overlap (starting from 33% of overlap between participants), brighter colors represent for up to 66% of overlap between participants.

Each row shows envelope mapping within each category (i.e. word, pseudoword, artificial and bird chirps), moving along the x-axis (from medial slices to lateral ones).

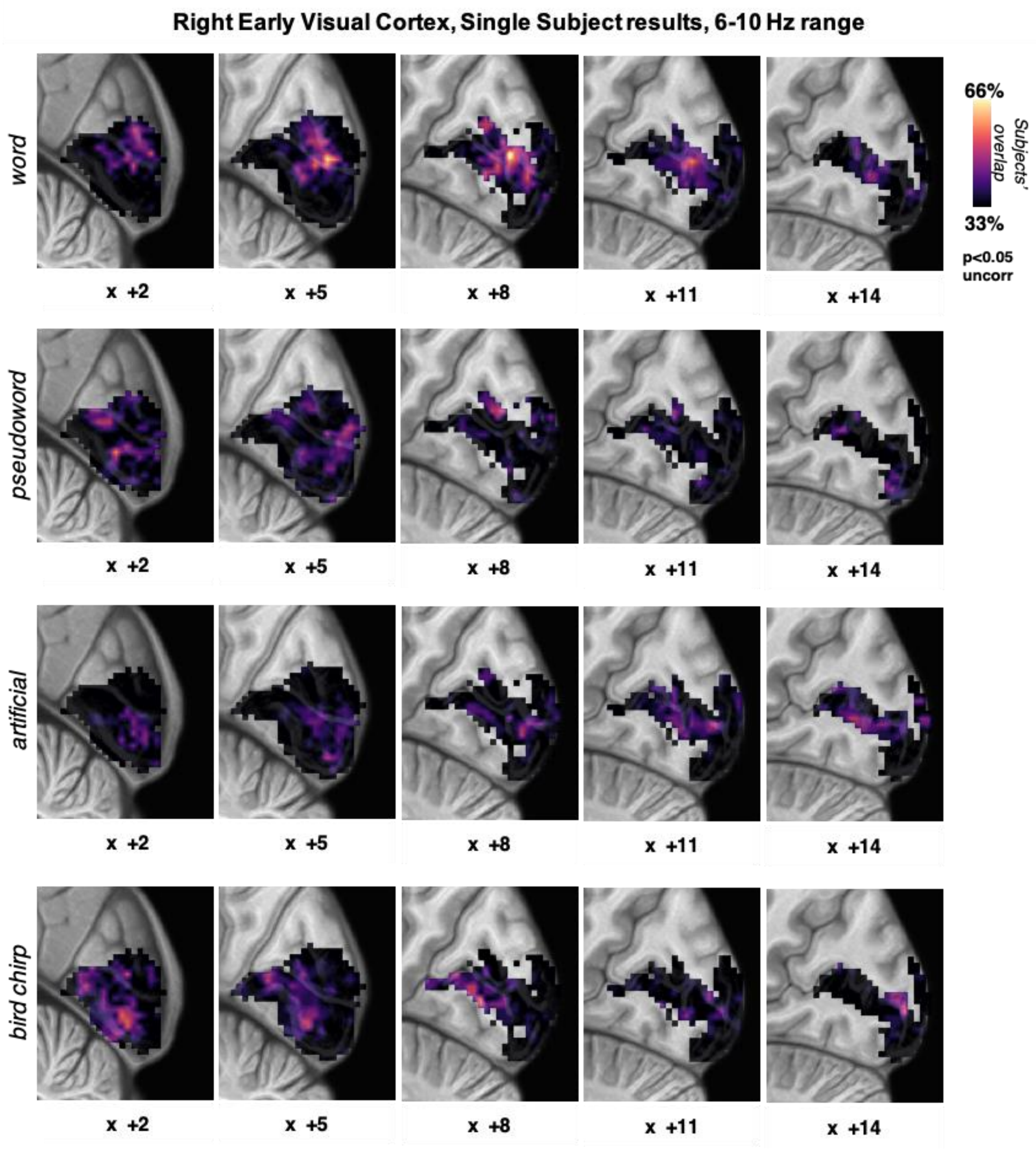


**Supplementary Figure 7. Envelope mapping single-subject results in the 6-10 Hz frequency range in right early visual cortex.** Within-category results of the single-subject envelope mapping in left Calcarine cortex are reported, representing the spatial overlap between participants (from 0 to 20 individuals; $p < 0.05$, uncorrected for multiple comparisons). Darker colors represent less overlap (starting from 33% of overlap between participants), brighter colors represent for up to 66% of overlap between participants.

Each row shows envelope mapping within each category (i.e. word, pseudoword, artificial and bird chirps), moving along the x-axis (from medial slices to lateral ones).

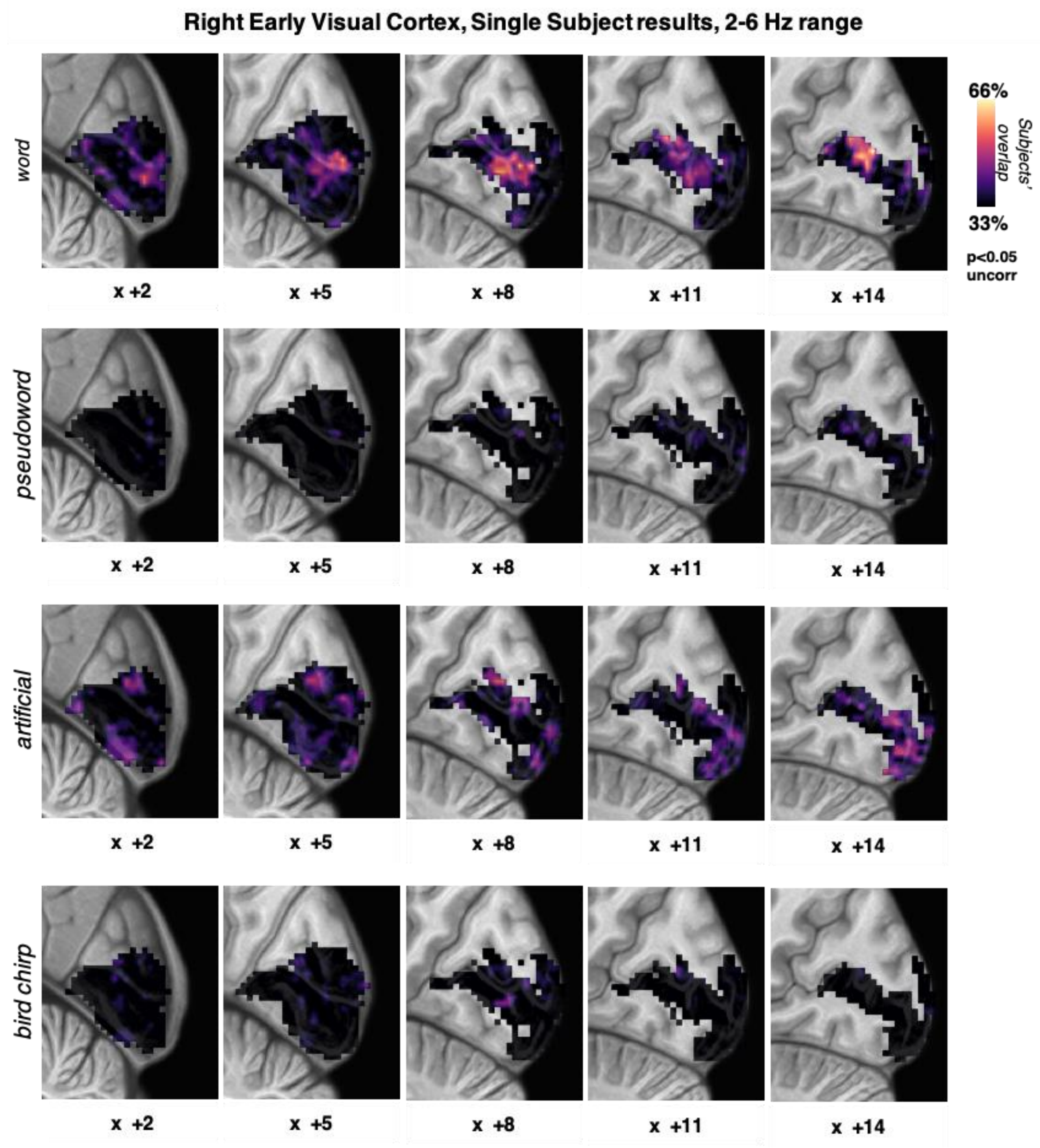


**Supplementary Figure 8. Envelope mapping single-subject results in the 2-6 Hz frequency range in right early visual cortex.** Within-category results of the single-subject envelope mapping in left Calcarine cortex are reported, representing the spatial overlap between participants (from 0 to 20 individuals; $p < 0.05$, uncorrected for multiple comparisons). Darker colors represent less overlap (starting from 33% of overlap between participants), brighter colors represent for up to 66% of overlap between participants.

Each row shows envelope mapping within each category (i.e. word, pseudoword, artificial and bird chirps), moving along the x-axis (from medial slices to lateral ones).

### Supplementary text

#### Comparisons between the mapping of Low and High frequency ranges

The number of significant voxels associated with High and Low-frequency ranges, averaged across all sound categories, did not differ in either left or right V1 (High vs. Low, left V1: $\chi2_{(1, 167)} = 0.203$, $p = 0.652$; right V1: $\chi2_{(1, 172)} = 2.852$, $p = 0.091$). In particular, in left V1 no difference between the number of voxels associated with High and Low-frequency ranges emerged for word ($\chi2_{(1, 167)} = 0.94$, $p = 0.759$) and pseudoword stimulus categories ($\chi2_{(1, 167)} = 0.114$, $p = 0.735$) in the left hemisphere. Similarly, no difference between the number of voxels associated with High and Low-frequency ranges emerged for the word category in the right hemisphere ($\chi2_{(1, 172)} = 1.381$, $p = 0.239$). No significant clusters of voxels were found in left V1 for both artificial sounds and birdsongs, nor in right V1 for artificial sounds. Selectively for the right V1 a greater mapping was found in response to bird chirps for the High compared to the Low-frequency range ($\chi2_{(1, 172)} = 16.78$, $p < 0.001$).

The number of voxels averaged across all sound categories, that in the Temporal ROI were associated with High frequency range, exceeded the ones with Low-frequency range, in both left and right hemispheres (left Temporal: $\chi2_{(1, 808)} = 5.434$, $p = 0.019$; right Temporal: $\chi2_{(1, 808)} = 18.046$, $p < 0.001$). In both the left and right hemispheres, the High frequency range was mapped in a greater number of voxels for pseudowords (left Temporal: $\chi2_{(1, 808)} = 8.090$, $p = 0.004$; right Temporal: $\chi2_{(1, 808)} = 7.205$, $p = 0.007$) and bird chirps (left Temporal: $\chi2_{(1, 808)} = 47.641$, $p < 0.001$; right Temporal: $\chi2_{(1, 808)} = 185.52$, $p < 0.001$). Only artificial sounds were associated with a greater mapping of Low compared with High-frequency range in the right hemisphere (right Temporal: $\chi2_{(1, 808)} = 20.55$, $p < 0.001$).
